# Polymer-directed inhibition of reversible to irreversible attachment prevents *Pseudomonas aeruginosa* biofilm formation

**DOI:** 10.1101/2022.01.08.475475

**Authors:** Alessandro M. Carabelli, Jean-Frédéric Dubern, Maria Papangeli, Nicola E. Farthing, Olutoba Sanni, Stephan Heeb, Andrew L. Hook, Morgan R. Alexander, Paul Williams

**Author notes:** Joint first authors (Alessandro M. Carabelli, Jean-Frédéric Dubern). Joint senior authors (Morgan R. Alexander, Paul Williams). Corresponding author: Biodiscovery Institute and School of Life Sciences, University of Nottingham, Nottingham, NG7 2RD, UK.

## Abstract

Non-toxic, biocompatible materials that inhibit bacterial biofilm formation on implanted medical devices and so prevent infection are urgently required. Weakly amphiphilic acrylate polymers with rigid hydrocarbon pendant groups resist bacterial biofilm formation *in vitro* and *in vivo* but the biological mechanism involved is not known. By comparing biofilm formation on polymers with the same acrylate backbone but with different pendant groups, we show that poly(ethylene glycol dicyclopentenyl ether acrylate; pEGdPEA) but not neopentyl glycol propoxylate diacrylate (pNGPDA) inhibited the transition from reversible to irreversible attachment. By using single-cell tracking algorithms and controlled flow microscopy we observed that fewer *Pseudomonas aeruginosa* PAO1 cells accumulated on pEGdPEA compared with pNGPDA. Bacteria reaching the pEGdPEA surface exhibited shorter residence times and greater asymmetric division with more cells departing from the surface post-cell division, characteristic of reversible attachment. Migrating cells on pEGdPEA deposited fewer exopolysaccharide trails and were unable top adhere strongly. Discrimination between the polymers required type IV pili and flagella. On pEGdPEA, the lack of accumulation of cyclic diguanylate or expression of *sadB* were consistent with the failure to transit from reversible to irreversible attachment. Constitutive expression of *sadB* increased surface adhesion sufficient to enable *P. aeruginosa* to form biofilms in a Mot flagellar stator dependent manner. These findings were extendable to other biofilm resistant acrylates highlighting their unique ability to inhibit reversible to irreversible attachment as a mechanism for preventing biofilm-associated infections.

**Significance:** Bacteria readily attach to surfaces forming biofilms. These are commonly associated with medical device-associated infections and highly refractory to antibiotics. Biocompatible, weakly amphiphilic acrylate polymers with large hydrophobic pendant groups that inhibit biofilm formation and can prevent such infections have been described. However, the biological mechanism involved is not understood. By comparing a biofilm-inhibiting with a biofilm-supporting acrylate, we showed that *Pseudomonas aeruginosa* PAO1 cells responded differentially to the two polymers and were unable to accumulate and adhere strongly, activate cyclic diguanylate signalling or transit from reversible to irreversible attachment on the inhibitory polymer. Constitutive expression of *sadB* increased surface adhesion sufficient to enable *P. aeruginosa* to form biofilms in a flagellar stator dependent manner overcoming the biofilm inhibitory properties of the polymer.

## Introduction

Implantable medical devices have enormously improved the quality of life for many millions of critically ill hospital patients (Percival, Suleman et al. 2015, VanEpps and Younger 2016). Such devices are highly susceptible to microbial colonization and device failure such that 50-70% of healthcare associated infections (HCAIs) can be directly attributable to biofilm formation. Despite improved hygiene measures taken to reduce bacterial contamination during implantation, the incidence of HCAIs is likely to grow as a consequence of increasing device utilization, population ageing, greater numbers of immunocompromised patients and the rise of multi-antibiotic resistant pathogens (Percival, Suleman et al. 2015, VanEpps and Younger 2016).

Biofilms are organized communities of bacteria embedded in a protective, self-generated extracellular matrix (ECM) that exhibit ‘emergent properties’ (Flemming, Wingender et al. 2016) that distinguish them from free-living cells such as their intrinsic tolerance to antibiotics and host immune defences and ability to cause chronic, persistent infections (Lebeaux, Ghigo et al. 2014, Tolker-Nielsen 2015, Flemming, Wingender et al. 2016, Ciofu and Tolker-Nielsen 2019). Effective strategies for the prevention, treatment and eradication of biofilm-associated infections are urgently required. Since biofilm formation is determined by the interplay between bacterial surface sensing, the prevailing environmental conditions and the physical and chemical properties of the device surface, the development of new preventive approaches depends on a better understanding of both the materials used to fabricate medical devices and the molecular and cellular mechanisms involved in biofilm initiation and development.

Most implanted medical devices are manufactured from synthetic polymers such as polypropylene, polyurethane and silicone (Greenhalgh, Dempsey-Hibbert et al. 2019). Historically these were selected largely for their biocompatibility, mechanical properties, ease of commercial production and low cost but with little consideration of susceptibility to biofilm-associated infections. Subsequent attempts to make such devices refractory to biofilm formation have predominantly focused on the incorporation of bactericidal agents (Greenhalgh, Dempsey-Hibbert et al. 2019). However, the main drawbacks of such a strategy are their limited efficacy due to active agent leaching, the tolerance of biofilms towards antimicrobial agents and selection for antibiotic resistance.

By screening a library of over 20,000 acrylate polymers for materials that inhibited biofilm formation (Hook, Chang et al. 2012, Hook, Chang et al. 2013), we discovered that while bacteria readily formed biofilms on almost all of the acrylates tested, we identified a small sub-set which inhibited biofilm development by multiple pathogens both *in vitro* and *in vivo* in mouse infection models (Hook, Chang et al. 2012, Hook, Chang et al. 2013). Using computational modelling combined with validation via in-house synthesis we derived a quantitative structure property relationship (QSPR). This demonstrated that biofilm inhibition required a weak amphiphilic acrylate backbone with a rigid hydrocarbon pendant group (Epa et al, 2014, Sanni, Chang et al. 2015, Mikulskis et al, 2018, Dundas, Sanni et al. 2019), a finding that could not have been predicted from current physico-chemical understanding of biofilm formation on diverse materials (Alexander and Williams 2017) (Carniello, Peterson et al. 2018). The first of this class of biofilm resistant polymers, poly(ethylene glycol dicyclopentenyl ether acrylate) (pEGdPEA; **Fig.1a**)) has recently been granted regulatory approval for clinical use as as a coating (BACTIGON^®^) for urinary tract catheters (Jeffery, Kalenderski et al. 2019).

Since pEGdPEA does not inhibit bacterial growth (Hook, Chang et al. 2012, Hook, Chang et al. 2013), it is likely to impact on the transition from the planktonic to a surface-attached lifestyle during the early stages of biofilm formation. To determine the biological basis for the biofilm inhibitory properties of pEGdPEA, we employed *Pseudomonas aeruginosa* as a model pathogen since it is commonly responsible for medical device-associated infections and has been intensively investigated with respect to biofilm development (Tolker-Nielsen 2015, O’Toole and Wong 2016, Valentini, Gonzalez et al. 2018). *P. aeruginosa* cells arriving from a bulk liquid weakly attach and then detach or stay and explore the surface by ‘walking’ or ‘crawling’. This depends on type IV pilus (T4P)-mediated twitching motility whereby the pilus extends and retracts to drag the bacterial cell body across the surface (Conrad, Gibiansky et al. 2011, O’Toole and Wong 2016). Transition to irreversible attachment, where cells orientate with their long axes parallel to the surface is the first committed step in biofilm formation (Petrova and Sauer 2012). This is followed by bacterial self-assembly into microcolonies and further development into mature biofilms where aggregated cells embedded in an extracellular matrix containing exopolysaccharides (EPS), proteins, lipids and extracellular DNA (eDNA) engage in physical and social interactions distinct from those of free-living bacterial cells (Petrova and Sauer 2012, Tolker-Nielsen 2015, Flemming, Wingender et al. 2016).

Transiting from a planktonic to a biofilm lifestyle depends on regulatory decisions relayed via sophisticated, integrated surface sensing networks that operate at the transcriptional, post-transcriptional and post-translational levels which in *P. aeruginosa* vary to some extent depending on the strain (Valentini and Filloux 2016, Chang 2018, Valentini, Gonzalez et al. 2018, Lee, Vachier et al. 2020). Flagella and T4P function not only in motility and as adhesins but also as surface mechanosensors activating cellular responses controlled by the intracellular second messengers, cyclic adenosine monophosphate (cAMP) and cyclic-di-guanylate (c-di-GMP) (O’Toole and Wong 2016). The latter controls the transition between motile and sessile lifestyles where elevated c-di-GMP levels promote biofilm formation but repress motility (Valentini and Filloux 2016). As bacterial cells reach a surface, mechanosensing through constraints on flagellar rotation serve to signal surface contact (Belas 2014) (Chaban, Hughes et al. 2015).

In *P. aeruginosa*, the increased c-di-GMP levels that arise during such interactions involve changes in flagellar stator engagement (Baker, Webster et al. 2019). Two different but exchangeable stator protein complexes (MotAB and MotCD) generate torque for flagellar rotation (Baker and O’Toole 2017) in which either stator complex is sufficient for swimming whereas both are required for mature biofilm formation (Toutain, Caizza et al. 2007). C-di-GMP levels regulate *P. aeruginosa* motility in part by influencing the localization of the two stators which in turn impacts on c-di-GMP production (Kuchma, Delalez et al. 2015, Baker, Webster et al. 2019). Surface sensing via the Pil-Chp complex and minor pilin PilY also causes c-di-GMP levels to rise rapidly triggering further T4P assembly via a mechanism that depends on MotAB and the c-diGMP receptor, FimW (Luo, Zhao et al. 2015) (Laventie, Sangermani et al. 2019).

During surface exploration and the early stages of biofim formation, *P. aeruginosa* produces an EPS such as Psl that acts as both an intercellular and a cell-surface ‘adhesive’ that is deposited in trails (Zhao, Tseng et al. 2013). These guide the T4P-mediated surface motility of visiting bacterial cells that secrete more Psl. This establishes a positive cycle by which accumulation of Psl promotes more intercellular adhesion and microcolony formation. Production of EPS is considered to be the dominant mechanism driving irreversible attachment of *P. aeruginosa* strain PAO1 lineages (Lee, Vachier et al. 2020).

To elucidate the biological basis by which this novel class of acrylate polymers inhibits bacterial biofilm formation we first followed the early stage interactions of single *P. aeruginosa* PAO1 cells with pEGdPEA compared with neopentyl glycol propoxylate diacrylate (pNGPDA; **Fig. 1b**) that has a more linear and flexible molecular structure but offers similar surface smoothness, stiffness, charge and hydrophobicity. We found that fewer cells accumulated on pEGdPEA, with those reaching the surface exhibiting shorter residence times, greater asymmetric division with more cells departing from the surface post-cell division. Using controlled flow microscopy, we determined that *P. aeruginosa* cells strongly adhered to pNGPDA but only weakly to pEGdPEA. Furthermore, on pEGPdEA, *P. aeruginosa* was unable to activate c-di-GMP signalling or to express *sadB* (PA5346), a gene required for the transition from reversible to irreversible attachment (Caiazza and O’Toole 2004, Caiazza, Merritt et al. 2007). By constitutively expressing *sadB*, surface adhesion could be increased sufficiently to enable *P. aeruginosa* form biofilms on pEGdPEA in a Mot stator dependent manner. These findings were extendable to other members of the pEGdPEA class of biofilm resistant acrylates demonstrating that inhibition of reversible to irreversible attachment is a conserved mechanism of action.

**Figure 1.**
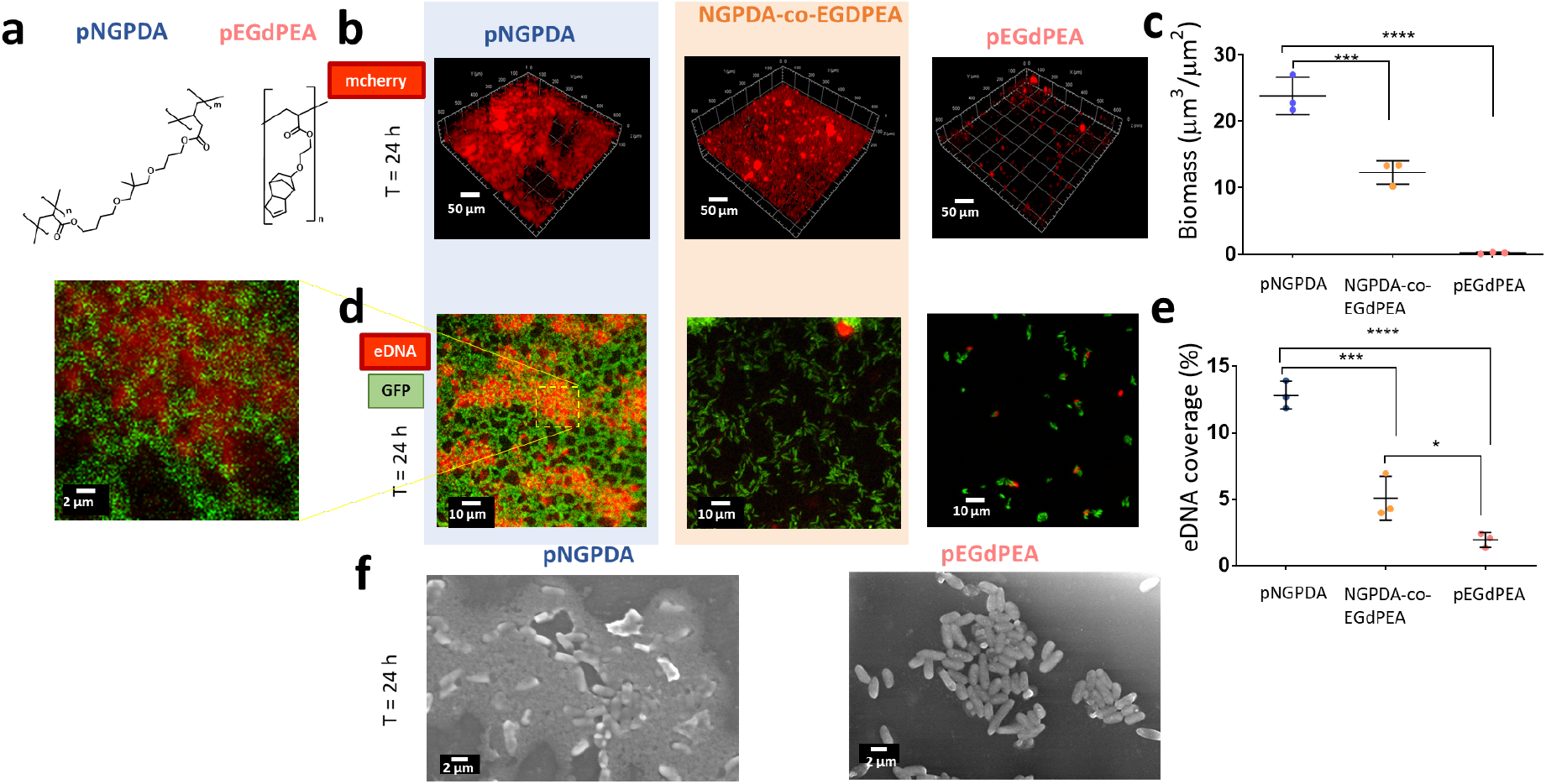
Mature biofilms form on pNGPDA but not on pEGDPEA (**a**) Chemical structures of NGPDA and EGdPEA monomers, (**b**) confocal images (10×, 0.3) of mCherry-tagged *P. aeruginosa* PAO1-W after 24 h incubation on pNGPDA (blue), pNGPDA-co-EGDPEA (orange), and pEGdPEA (pink). Scale bar, 50 µm; (**c**) Quantification of bacterial biomass on the 3 polymer surfaces; (**d**) confocal images (40×,1.2) of GFP-tagged *P. aeruginosa* PAO1-W after 24 h on pNGPDA (left), pNGPDA-co-EGDPEA (middle), and pEGdPEA (right) and stained for eDNA with propidium bromide and (**e**) quantification of eDNA coverage in (**d**) eDNA was stained through propidium iodide (red). Left-hand panel shows higher magnification of biofilm on pNGPDA; (**f**) representative ESEM images of PAO1-W grown for 24 h on pNGPDA (left) and pEGdPEA (right). Images captured at 10,000×. Scale bar, 2 µ m. Error bars show ± 1 standard deviation (N=3). Significance was determined by analysis of variance One-way ANOVA and Tukey’s post-test comparison for differences between the indicated samples. **** *p*<0.001, *** *p*<0.001, * *p*<0.05.

## Results

### *P. aeruginosa* does not form mature biofilms on pEGDPEA

Pro- and anti-biofilm acrylate polymer coatings of pNGPDA and pEGdPEA respectively (**Fig. 1a**) together with a 1:1 NGPDA-EGdPEA copolymer were prepared by UV photoinitiated radical polymerisation on glass coverslips in a reduced oxygen environment (**Table S1**). Surface chemical (TOF-SIMS) and physical (wettability, surface charge and roughness) properties measurements (**Figs. S1-3**) confirmed the chemical identity of the polymer surfaces and confirmed homogenous coverage distribution of NGPDA and EGdPEA within the copolymer. The surfaces of both pNGPDA and pEGdPEA were similarly negatively charged (-52.6 ± 2.55 mV and -64.6 ± 1.25 mV respectively in RPMI). The water contact angles for pEGdPEA (86.3 ± 2.2°) and pNGPDA (69.8 ± 2.7°) were similar.

*P. aeruginosa* PAO1 (Washington collection sub-line; PAO1-W) biofilm development on pEGdPEA, pNGPDA and the NGPDA-EGdPEA copolymer coated onto borosilicate glass coverslips was quantified using confocal microscopy. Representative images after incubation for 24 h in RPMI-1640 are shown in **Fig.1b** and the biomass quantified (**Fig. 1c**). Bacterial biomass was reduced from 23.8 ± 2.8 µm^3^ µm^-2^ on NPGDA to 0.2 ± 0.1 µm^3^ µm^-2^ on pEGdPEA. An intermediate biomass was measured on p(NGPDA-co-EGdPEA) (12.3 ± 1.8 µm^3^ µm^-2^) compared with pEGdPEA. After 72 h (**Fig. S4**). *P. aeruginosa* biofilm on pEGdPEA and p(NGPDA-co-EGdPEA) (1:1) was 0.4 ± 0.2 µm^3^ µm^-2^ and 0.7 ± 0.1 and µm^3^ µm^-2^ respectively, whereas pNGPDA exhibited much higher biofilm biomass (47.8 ± 7.9 µm^3^ µm^-2^). Biofilm on pNGPDA also exhibited higher levels of eDNA consistent with the formation of mature biofilms in contrast to p(NGPDA-co-EGdPEA) and pEGdPEA (**Fig. 1d-e**). Environmental scanning electron microscopy (ESEM) (**Fig. 1f**) confirmed the presence of bacteria embedded in an extensive ECM on pNGPDA whereas in contrast, on pEGdPEA, only a few *P. aeruginosa* cell clusters lacking an ECM were observed.

### *P. aeruginosa* cells show differential early stage interactions with with pEGdPEA and pNGPDA

To explore the early stage interactions between single bacterial cells and the polymers, we employed a multimode microscope (Hook, Flewellen et al. 2019) that enabled observation of individual bacterial cells at the polymer surface and within the 3D volume above the polymer surfaces on coated coverslips in a custom-made holder. Since the near surface runs of *P. aeruginosa* differ from those in bulk liquid (Hook, Flewellen et al. 2019), we compared the 3D trajectories of near-surface bacteria swimming in the bulk liquid up to 50 µm above each surface using digital holographic microscopy (DHM). No differences in the speed of cells swimming above pNGPDA (62.1± 11.9 μm/s) and pEGdPEA (61.5±11.7) during the first 10 min were noted (**Fig. S5a)**. To observe bacterial behaviour on the surface, differential interference (DIC) microscopy images were acquired at 0.5 Hz over the 3 h period immediately after inoculation from the near surface region of each polymer to observe bacterial behaviour at the surface ( ). Over the first 60 min, similar numbers of surface-associated cells were present on pNGPDA and pEGdPEA (**Fig. S5b** and **c**), suggesting that neither polymer prevented the reversible attachment of bacterial cells. However, after 3 h, more bacteria accumulated on pNGPDA than on pEGdPEA (*p*<0.01; unpaired t-test analysis; **Fig. S5d** and **e**).

The number of cells on a surface change either as a consequence of cells arriving, leaving or dividing. Cell doubling times and surface residence on each polymer surface between t = 1 h and t = 2 h were determined using a cell tracker algorithm (Hook, Flewellen et al. 2019). *P. aeruginosa* replicated with similar doubling times (59 ± 8 min and 58 ± 6 (N=3, n=20)) on both pNGPDA and pEGdPEA respectively, but were observed to have longer and more variable residence times on pNGPDA (**Fig. 2a**). To explore the post-division fate of surface-associated cells, we investigated whether both daughter cells detached (two-legged branching), only one detached (one-legged branching) or both remained (zero-legged branching) on the surface (**Fig. 2b**) (Lee, Vachier et al. 2020) (Lee, de Anda et al. 2018). As shown in **Fig. 2c**, zero-legged branching was significantly (*p*<0.001) greater on pNGPDA (83 ± 1%) compared with pEGdPEA (53 ± 10%), whereas one-legged branching was significantly greater on pEGdPEA (44 ± 7%) than on pNGPDA (17 ± 1%) (*p*<0.001). Therefore, the change in surface coverage observed on the two polymers after 3 h is likely to be due to differences in the rates of departure after cell division.

**Figure 2.**
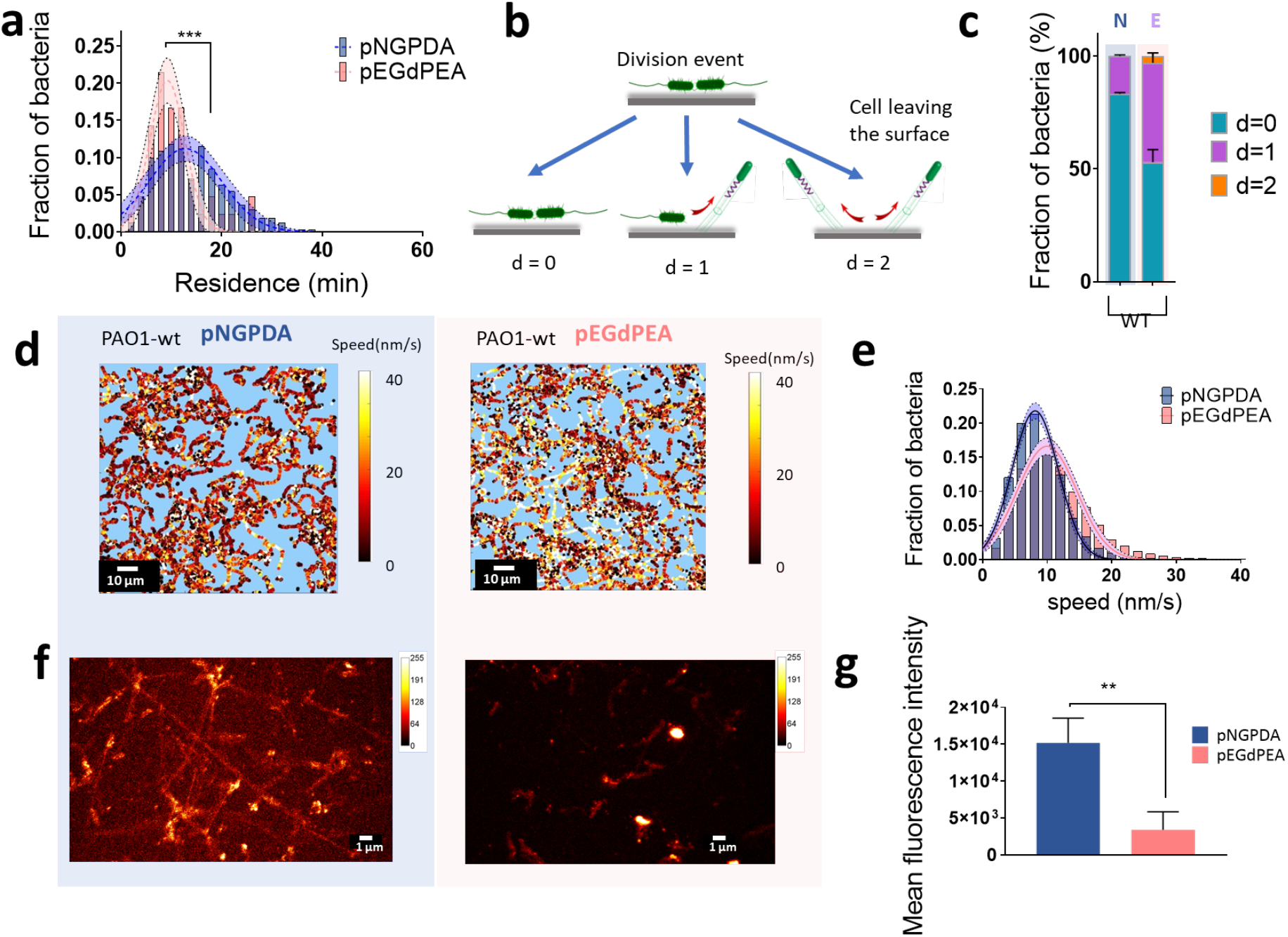
Single cell tracking of *P. aeruginosa* migration on, and departure from, pNGPDA and pEGDPEA. (**a**) Residence time of PAO1-W on pNGPDA (blue bar) and pEGdPEA (pink bar) between t = 1 h to t = 2 h after inoculation. Solid bars represent fraction of bacteria at different residence times (bin = 2 min). Dashed lines represent data fit to Gaussian functions for pNGPDA (blue) and pEGdPEA (pink). Shaded areas represent 95% CI of the fit deviation. R^2^=0.86 for pNGPDA and R^2^=0.91 for pEGdPEA (N=3 considering more than 700 bacteria). Welch’s t test was applied to compare distribution assuming different variance. *** *p*<0.001; (**b**) Schematic of the branching division events on a surface; (**c**) Proportion of the population for PAO1-W in different branching divisions where pNGPDA (blue) and pEGdPEA (pink). Values obtained considering n>100 doubling events (N=3) between t = 1 h to t = 2 h from inoculation. PAO1-W branching distribution for the two polymers was found to be different (*p*<0.001) through two-way ANOVA and Sidak’s multiple comparison; (**d**) Representative colour maps of PAO1-W twitching tracks on pNGPDA and pEGdPEA measured over 1 h of exposure to surfaces. Colours show different speeds as indicated by the intensity scale. Scale bar, 10 µm; (**e**) Histogram of PAO1-W speeds on pNGPDA and pEGdPEA (N=3). Bin = 2. Dashed lines represent data fit to Gaussian functions. Shaded areas represent 95% CI of the fits. **** *p*<0.0001 using Welch’s t test assuming different variance; (**f**) Confocal fluorescent microscope images captured at t = 2.5 h on pEGdPEA (right) and pNGPDA (left), colour map shows fluorescence levels, Scale bars, 10 µm; (**g**) quantification of ConA-Alexa 633 bound to EPS on each polymer surface. Scale bars, 10 µm. Unpaired t-test (\*\**p*<0.01). Error bars show ± 1 standard deviation (N=3).

In addition to swimming in the medium above and near the surface *P. aeruginosa* cells can migrate over surfaces by deploying T4P-mediated twitching motility to ‘walk’ (oriented perpendicular to the surface) or ‘crawl’ (parallel to the surface) at speeds in the region of 5-80 nm/s ((Conrad, Gibiansky et al. 2011) (O’Toole and Wong 2016) (Rodesney, Roman et al. 2017). Hence, we explored the twitching directionality and speed on both polymers. **Fig. 2d** shows examples of bacterial cell tracking on the two polymer surfaces where instantaneous speed within the tracks is indicated using a heat scale. Directionality was evaluated by considering the radius of curvature (**Fig. S6a**), whereby higher values of this parameter correspond to more linear trajectories (Utada, Bennett et al. 2014). The fraction of twitching cells (55 ± 16% and 58 ± 3% on pNGPDA and pEGdPEA respectively), their directionality (**Fig S6a**) and mean twitching speeds (9 ± 4 nm/s and 11 ± 6 nm/s (**Fig 2e**) on the two polymers were similar. However on pEGdPEA a small population of faster migrating cells is apparent (**Fig. 2e**). The proportion of cells as a function of tilting fraction (where 0 = flat and 1 = tilting up) appears to be greater on pEGdPEA than on pNGPDA (**Fig. S6b** and **c**).

During the early stages of biofilm development as *P. aeruginosa* PAO1 cells move over surfaces, they lay down tracks containing the mannose-rich EPS adhesive, Psl (Zhao, Tseng et al. 2013). Using DIC and confocal laser scanning microscopy to track cells over 2.5 h post inoculation, and by staining for EPS with ConA-Alexa 633, we observed that *P. aeruginosa* lays down EPS trails on pNGPDA but not on pEGdPEA (**Fig. 2f** and **g**).

The differential colonisation and daughter cell fates on the two polymer surfaces suggested a difference in the bacteria-polymer interaction. To investigate this, the strength of cell adhesion to each polymer was measured using a shear stress detachment assay in a microfluidics chamber 10 min after inoculation. This apparatus allowed application of gradually increasing flow rates designed to exert a shear stress *τ* from 0.6 up to 630.0 Pa over a period of 4 min, whilst simultaneous image capture followed the removal of cells from the surface. Subsequent image analysis to capture cell number as a function of time, allowed a shear stress, *τ*_50_ (the shear stress required to remove 50% of attached cells), to be measured (Arpa-Sancet, Christophis et al. 2012). The *τ*_5_ was was significantly (*p*<0.0001) reduced from 87.6 ± 15.0 Pa on pNGPDA to 3.2 ± 0.8 Pa for bacteria grown on pEGdPEA (**Fig. S7**). An intermediate *τ*_50_ of 31.2 ± 15.0 Pa was observed for the NGPDA-co-EGdPEA polymer.

### Are flagella and T4P required for biofilm formation on pNGPDA and pEGdPEA?

Biofilm formation was quantified after 24 h growth comparing wild type *P. aeruginosa* T4P (Δ*pilA*), flagellin (Δ*fliC*), Δ*pilA*Δ*fliC* and flagella stator (Δ*motAB,* Δ*motCD* and Δ*motABCD*) deletion mutants (**Fig. 3 a** and **b**). The Δ*pilA* and Δ*pilA*Δ*fliC* were unable to form biofilms on either polymer whereas the Δ*fliC* mutant, which lacks a flagellum, formed biofilm on pNGPDA although it was ∼46% less than the wild type. Neither strain formed a biofilm on pEGdPEA. Therefore, bacteria lacking flagella still differentiate between the two surfaces. *P. aeruginosa* Δ*motAB* and Δ*motCD* stator mutants both produce flagella and swim normally whereas the Δ*motABCD* mutant has paralyzed flagella and is non-motile (Doyle et al, 2004; Hook, Flewellen et al. 2019). The *mot* stator mutants produced relatively little biofilm on pNGPDA compared with the wild-type. However the Δ*motCD* mutant (19.0 ± 3.4 µm^3^/µm^2^) produced statistically significant greater biomass (*p*<0.001) on pEGdPEA than the wild type (6.0 ± 3.4 µm^3^/µm^2^). (**Fig. 3a** and **b**). These data indicate that both T4P and the *mot* stators are essential for mature biofilm formation on both polymers whereas the *fliC* mutant remains capable of forming a biofilm on pNGPDA albeit with a reduced biomass compared with wild type.

**Figure 3.**
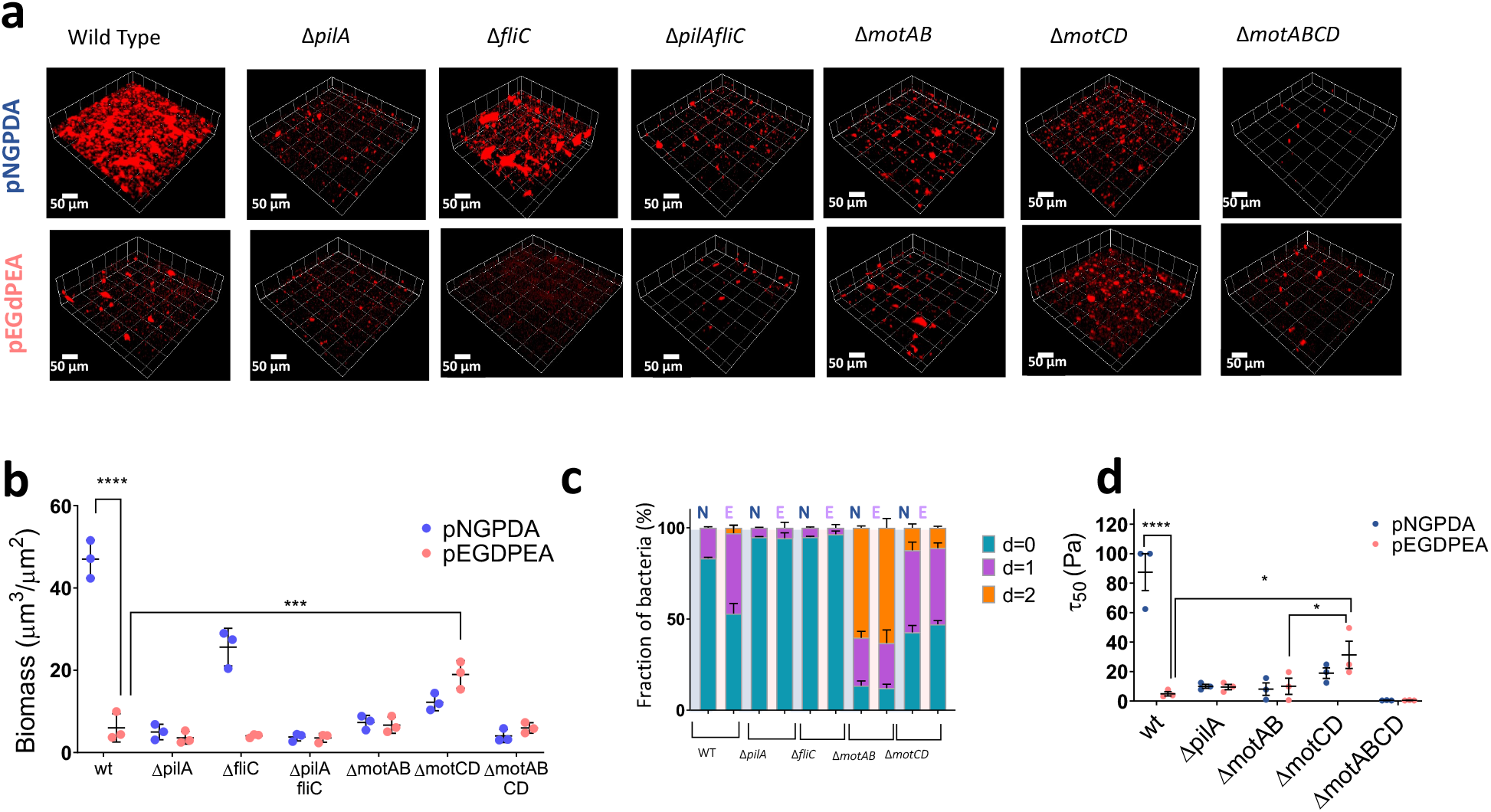
Differential behavior of *P. aeruginosa* PAO1-W wild type, flagellar (*fliC*), T4P (*pilA*) and *mot* stator mutants on pNGPDA and pEGDPEA. (**a**) Confocal images (10×, 0.3) of *P. aeruginosa m-cherry* tagged PAO1-W, Δ*pilA*, Δ*fliC,* Δ*pilAfliC* Δ*motAB*, Δ*motCD and* Δ*motABCD* after 24 h on pNGPDA (top) and pEGdPEA (bottom). Scale bar, 50 µm. (**b**) Quantification of *P. aeruginosa* wild type, Δ*pilA*, Δ*motAB*, Δ*motCD* and Δ*motABCD* biofilm biomass after 24 h on pNGPDA and pEGdPEA. Samples were washed 2× in PBS and 1× in H_2_O. Scale bar is 50 µm . Significance was determined by 2-way ANOVA multiple comparison with Sidak’s correction for differences between the indicated samples. \*\*\*\**p*<0.0001, \*\*\**p*<0.001 and \**p*<0.05. (**c**) Proportion of the population in each branching divisions post cell division on pNGPDA (N, blue) and pEGdPEA (E, pink). Values obtained considering n>100 doubling events for 3 independent replicates (N=3) between t = 1 h and t = 2 h after inoculation. (**d**) Shear stress τ_50_ of PAO1-W, Δ *pilA*, Δ *motAB*, Δ*motCD* and Δ*motABCD* cells on pEGdPEA (pink) and pNGPDA (blue).

To further elucidate the role of *P. aeruginosa* flagella and T4P on surface residence and detachment events, we determined the post-division fate of the Δ*pilA*, Δ*fliC*, Δ*pilA*Δ*fliC*, Δ*motAB* and Δ*motCD* mutants on the two polymers. A comparison of early stage (t = 1 to 2 h) surface-associated wild-type *P. aeruginosa* cells with Δ*pilA* and Δ*fliC* mutants post cell division revealed that, in contrast to the wild-type, both mutants showed predominantly zero-legged branching (d=0). This is consistent with a requirement for both T4P and flagella for “active” detachment (**Fig. 3c**)(Conrad, Gibiansky et al. 2011). In contrast, Δ*motAB* cells showed a greater propensity (d=2) for leaving both polymer surfaces compared with wild-type with 36 ± 5% and 24 ± 11% of the daughter cells leaving pNGPDA and pEGdPEA respectively. The Δ*motCD* mutant behaved similarly on both polymer surfaces but differed from the Δ*motAB* mutant with a higher proportion of d=0 cells and resembled that of the wild-type on pEGdPEA with most cells showing one-legged branching (**Fig. 3c** and **Table S2**). Thus loss of either stator reduced the differential behaviour of *P. aeruginosa* on both polymers.

To investigate further the role of flagella and T4P on *P. aeruginosa* early stage surface attachment, we determined the relative strengths of adhesion of the Δ*pilA* and Δ*fliC* mutants compared with the wild type (**Fig. 3d** and **Fig. S7**). The shear stress (*τ*_50_) required to detach the Δ*pilA* mutant was similar on both pNGPDA and pEGdPEA (10.0 ± 2.3 and 9.4 ± 3.1 Pa respectively) compared with 87.6 ± 15.0 Pa and 3.2 ± 0.8 Pa respectively for the wild-type. These data demonstrate that T4P are required for surface adhesion to pNGPDA, but are not able to adhere strongly to pEGdPEA (**Fig. 3d**). Surprisingly, despite the reduction in biofilm observed on pNGPDA compared with wild type (**Fig.3 b**) the Δ*fliC* mutant adhered very strongly to both pNGPDA and EGdPEA and could not be detached from either polymer at the highest level of shear stress that could be applied (**Fig. S7c**). To determine whether the flagellar stators influenced adhesion to pNGPDA and pEGdPEA, the *mot* mutants were also tested in the microfluidic detachment assay (**Fig. 3d**). The adhesion strength of the *motAB* and *motCD* mutants on pNGPDA was substantially reduced compared with the wild-type with *τ*_50_ values of 8.2 ± 7.3 and 18.9 ± 6.2 Pa respectively and was significantly further reduced for the non-motile *motABCD* mutant (*τ*_50_ = 0.5 ± 0.1 Pa on both polymers) (**Fig. 3d**). On pEGdPEA, although the *motAB* mutant presented similar attachment strength to the wild-type (10.0 ± 9.5 Pa), the *motCD* mutant exhibited increased adherence (31.3 ± 16.0 Pa; *p<0.05* in comparison to the wild-type). These data show that T4P and the flagellar stators are required for *P. aeruginosa* adhesion and hence biofilm formation on pNGPDA but were unable to facilitate strong adhesion to pEGdPEA.

### *P. aeruginosa* is Unable to Progress from Reversible to Irreversible Attachment on pEGdPEA

The second messenger c-di-GMP plays a central role in determining the transition from a motile to a sessile lifestyle and impacts on flagellar- and TP4-mediated motility as well as exopolysaccharide production (Valentini and Filloux 2016, Valentini, Gonzalez et al. 2018). To understand how interactions with pNGPDA and pEGdPEA impact on c-di-GMP, we employed a *cdrA::gfp* fusion as a surrogate reporter for c-di-GMP (Rybtke, Borlee et al. 2012) that can be followed in single cells using time-lapse epifluorescence microscopy. On pNGPDA, the fluorescence intensity of the *cdrA* promoter fusion strongly increased over time as cells accumulated on the polymer surface over 3 h. This contrasted with the data obtained on pEGdPEA, where 30% lower fluorescence intensity (*p<0.*0001) was observed consistent with lower levels of c-di-GMP production (*F* values were 46.3 ± 1.8 and 65.6 ± 1.0 on pEGdPEA and pNGPDA respectively) (**Fig. 4a-b**). By plotting the fluorescence of each individual cell 3 h after inoculation, we observed individual cells with low and high c-di-GMP reporter activity on pNGPDA (**Fig. 4b**) indicating more heterogenous *cdrA* promoter expression than on pEGdPEA.

**Figure 4.**
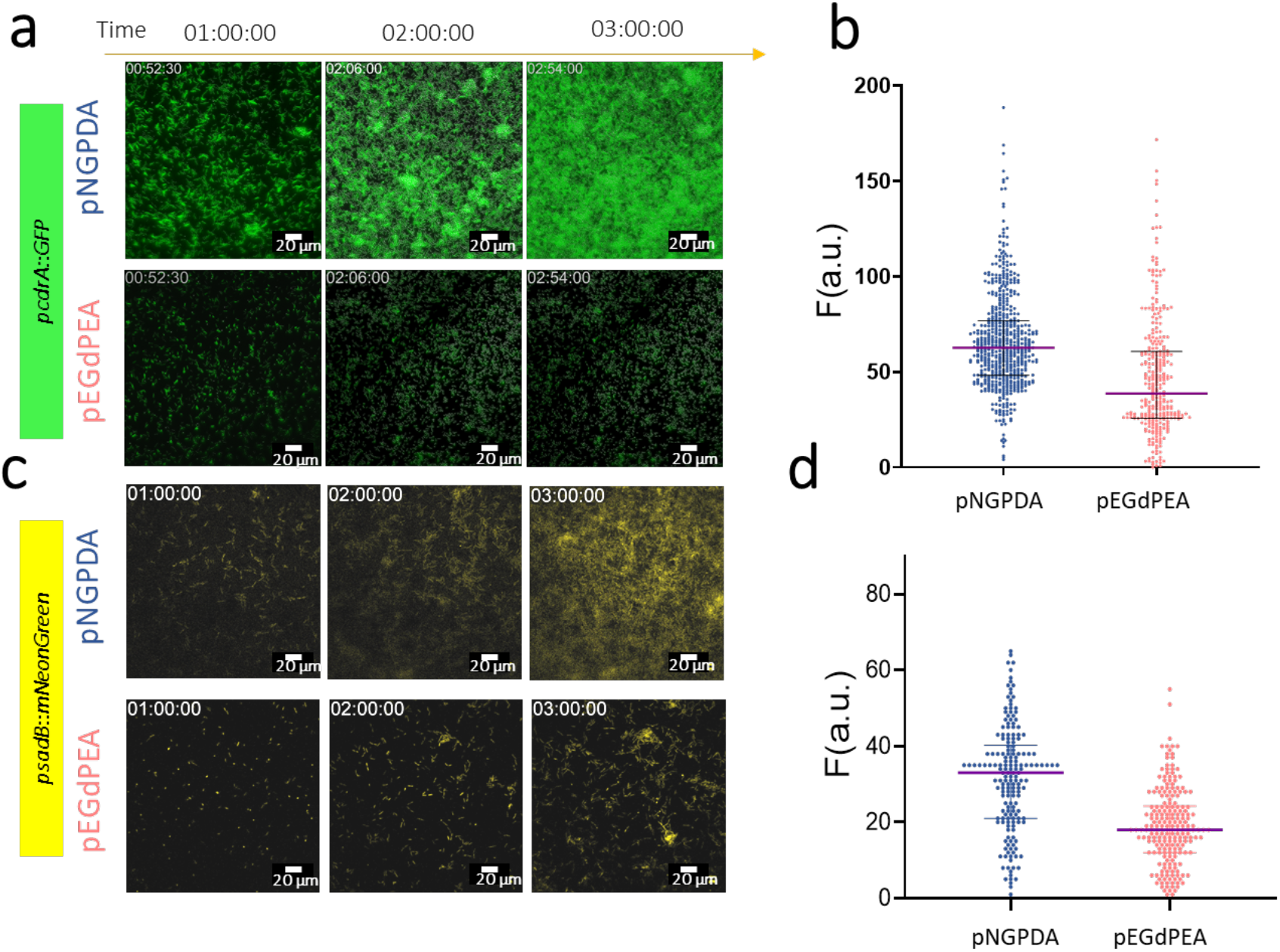
The expression of *cdrA* (**a** and **b**) and *sadB* (**c** and **d**) promoters over time in *P. aeruginosa* on pNGPDA and pEGDPEA. (**a**) Representative epifluorescence microscopy time series images of *P. aeruginosa* PAO1-W transformed with pCdrA’::*gfp*(ASV) and (**b**) single cell quantification at t = 3 h on pNGPDA (blue) and pEGdPEA (pink); (**c**) Representative epifluorescence microscopy time series images of *P. aeruginosa* pSadB’::*mNeonGreen* and (**d**) quantification at t = 3 h on pNGPDA (blue) and pEGdPEA (pink) Error bars show ± 1 standard deviation (N=3). Scale bars, 20 µm.

During the early stages of biofilm formation, the reversible attachment of *P. aeruginosa* cells to a surface becomes irreversible through the co-ordinated activation of multiple pathways (Petrova and Sauer 2012).(Petrova and Sauer 2012) One of these involves the *sadB* gene product (O’Toole and Kolter 1998) (O’Toole and Kolter 1998, Caiazza and O’Toole 2004, Caiazza, Merritt et al. 2007). We therefore hypothesized that *P. aeruginosa* cells may be unable to undergo reversible to irreversible attachment to pEGdPEA as they are unable to induce sadB expression. To explore this possibility we constructed a p*sadB*’*::mNeongreen* promoter fusion and followed *sadB* expression over time by time-lapse epifluorescence microscopy. Higher levels of fluorescence were observed after 3 h incubation on pNGPDA compared with pEGdPEA consistent with differential expression of *sadB* (**Fig. 4c**) In addition, quantitative single cell analysis, showed that *sadB* expression on pNGPDA was more heterogeneous than on pEGdPEA (**Fig. 4d**) similar to that noted for the *cdrA::gfp* fusion.

Since these data are consistent with the inability of *P. aeruginosa* to induce *sadB* on pEGdPEA, we constructed a *sadB* constitutive expression vector incorporating a non-native (ptac) promoter and a *sadB* deletion mutant. **Fig. 5a** shows representative confocal images of biofilm formation on pNGPDA and pEGdPEA by the wild-type (transformed with the empty plasmid vector), Δ*sadB* deletion mutant, the complemented Δ*sadB* mutant and wildtype constitutively expressing *sadB.* The biofilm biomass of *P. aeruginosa* pME6032 reduced from 9.8 ± 1.2 µm^3^ µm^-2^ on pNPGDA to 0.9 ± 0.3 µm^3^ µm^-2^ on pEGdPEA consistent with our previous data. While the Δ*sadB* mutant was unable to form biofilm on either surface, constitutive expression of *sadB* in the wild-type enabled *P. aeruginosa* to form a biofilm on pEGdPEA (14.5 ± 0.7 µm^3^ µm^-2^) (**Fig. 5a-b**). Similarly, genetic complementation of the Δ*sadB* mutant restored biofilm on both polymer surfaces. Furthermore, shear stress measurements revealed that the shear force required to detach the wild-type cells constitutively expressing *sadB* cells from pEGdPEA was *τ*_50_ = 31.3 ± 16.0 Pa. This is significantly greater than the wild-type (\*\**p*<0.01). The weakest adhesion was observed for the *sadB* mutant (0.5 ± 0.1 Pa) (**Fig. 5e**).

**Figure 5.**
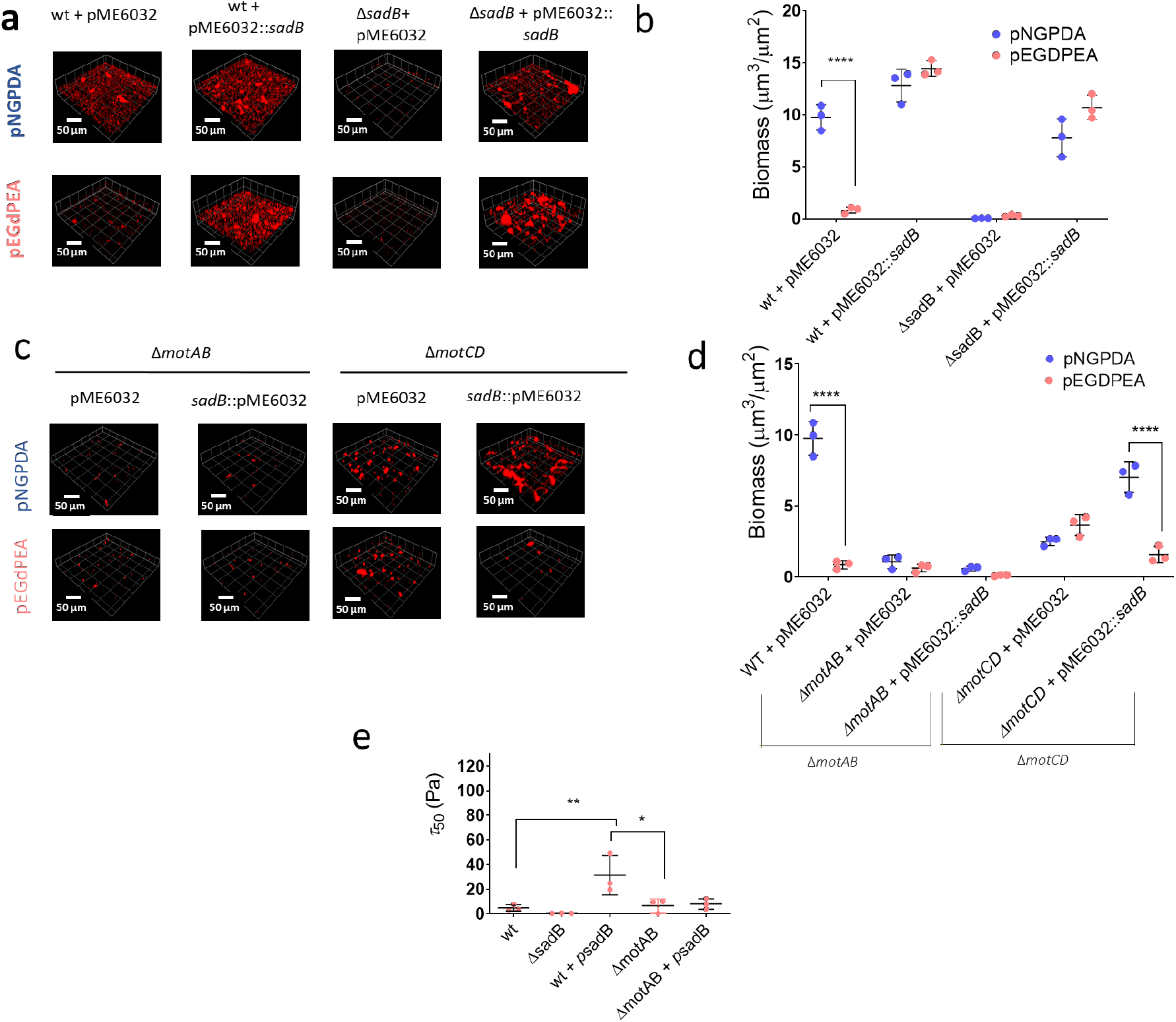
Constitutive *sadB* overexpression increases adhesion strength and overcomes the biofilm resistance of pEGDPEA to *P. aeruginosa* PAO1-W in a stator dependent manner. (**a**) Confocal images (10×, 0.3) and quantification (**b**) of mCherry tagged *P. aeruginosa* PAO1-W pME6032 (empty vector), PAO1-W pME6032::*sadB*, Δ *sadB* pME6032 (empty vector) and Δ*sadB* pME6032::*sadB* after 24 h on pNGPDA (top) and pEGdPEA (bottom); (**c**) Confocal images (10×, 0.3) and quantification (**d**) of m-cherry tagged *P. aeruginosa* PAO1 Δ*motAB* and Δ*motCD* transformed with pME6032 (empty vector) or pME6032::*sadB* after 24 h on pNGPDA (top) and pEGdPEA (bottom). (**d**) Shear stress (τ_50_) necessary to detach 50% of PAO1-W wild-type or Δ*motAB* and Δ*motCD*. (e) PAO1-W pME6032::*sadB*, Δ*sadB* pME6032 (empty vector) cells on pEGdPEA. Error bars represent ± 1 standard deviation unit (N=3). (**e**) Shear stress (τ_50_) necessary to detach 50% of PAO1-W, PAO1-W pME6032::*sadB*, Δ*sadB,* Δ*motAB* or Δ*motAB* pME6032::*sadB* cells on pEGdPEA. Error bars represent ± 1 standard deviation unit (N=3). Significance was determined by Anova analysis using Tuckey’s multiple comparison for differences between samples. \*\**p*<0.01 and \**p*<0.05. Scale bars in (**a**) and (**c**), 50 µm.

To determine whether biofilm formation on pEGdPEA as a result of the constitutive expression of *sadB* was dependent on one or both of the *mot* stators, we quantified biofilm formation on both pNGPDA and pEGdPEA by the *motAB* and *motCD* mutants constitutively expressing *sadB*. **Fig. 5c and d** shows that constitutive expression of *sadB* in the Δ*motAB* mutant was unable to restore biofilm formation on either polymer. However, a statistically significant increase in biofilm formation was observed on pNGPDA but not pEGdPEA for the Δ*motCD* mutant expressing *sadB*. Consistent with these findings, we clearly observed that constitutive over-expression of *sadB* in the wildtype but not in the *motABCD* stator mutant on pEGdPEA restored cyclic diguanylate production as reported by the *cdrA’::gfp* fusion (**Fig S8**).

### Constitutive expression of *sadB* restores the ability of *P. aeruginosa* to form biofilms on chemically related biofilm-resistant acrylates

To determine whether constitutive expression of *sadB* could also overcome the biofilm resistance of other amphiphilic acrylates related to pEGdPEA that fulfil the QSPR requirements for biofilm inhibition (Epa et al, 2014; Sanni, Chang et al. 2015, Mikulskis et al, 2018, Dundas, Sanni et al. 2019), we selected three polymers with pendant groups distinct from the dicyclopentenyl ether of pEGdPEA but that had also been previously shown to prevent bacterial biofilm formation (Hook, Chang et al. 2013). These were benzyl methacrylate (BnMA), isobornyl methacrylate (iBMA) and trimethylcyclohexyl methacrylate (TMCHMA). **Fig. 6a-b** shows that significant (p<0.0001) increases in biofilm formation by *P. aeruginosa* constitutively expressing *sadB* were apparent for all three biofilm inhibitory polymers compared with the vector control.

**Figure 6.**
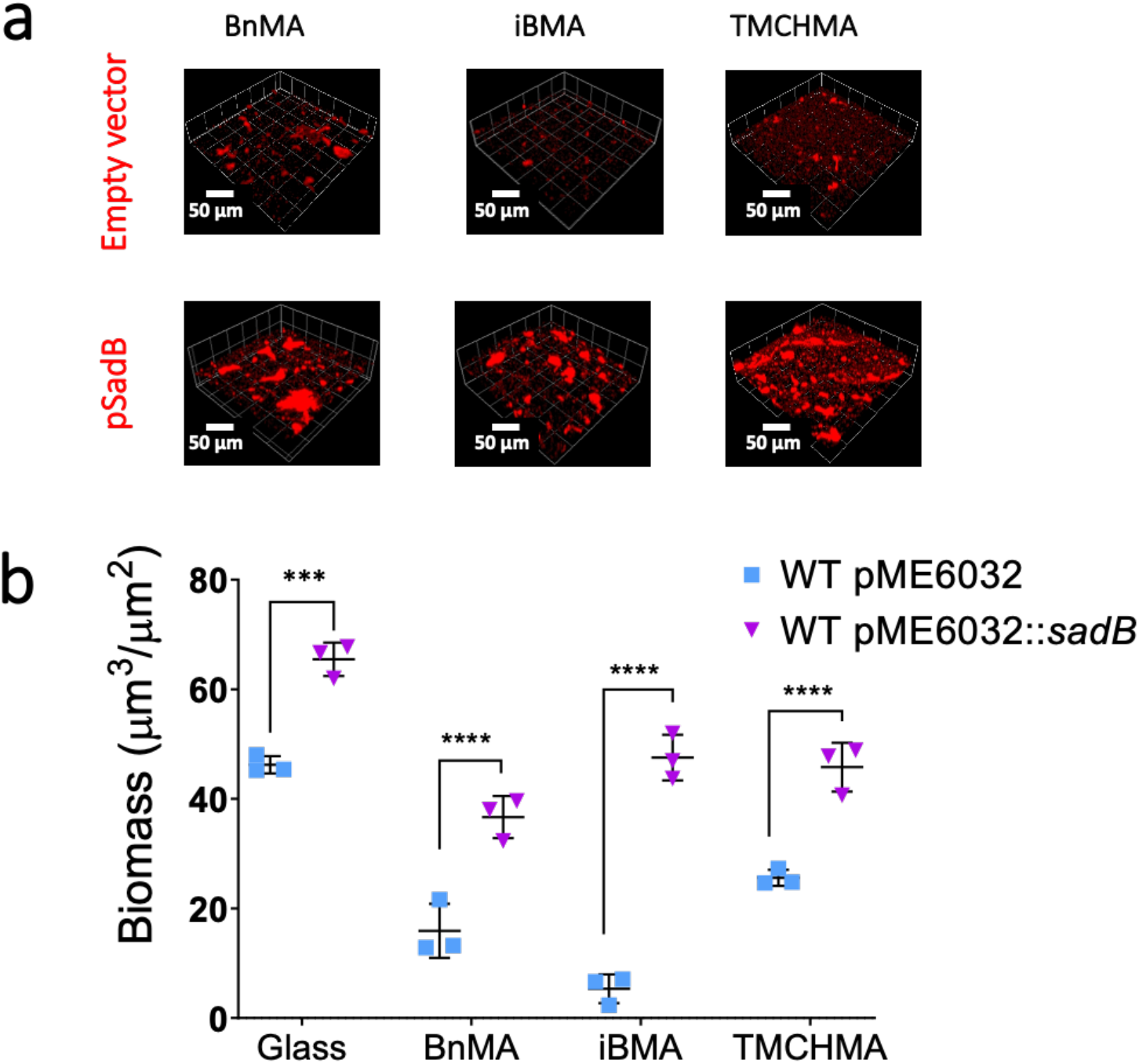
Constitutive expression of *sadB* in *P. aeruginosa* PAO1-W overcomes the biofilm resistance of *P. aeruginosa* PAO1-W on biofilm inhibiting acrylate polymers. (**a**) Confocal images (10×, 0.3) and quantification (**b**) of mCherry tagged *P. aeruginosa* PAO1-W pME6032 (empty vector, blue squares) PAO1-W pME6032::*sadB* (violet triangles), after 24 h on BnMA, iBMA and TMCHMA. Scale bar, 50 µ m. Error bars show ± 1 standard deviation (N=3).

## Discussion

To begin unravelling the mechanism by which specific acrylate polymers inhibited bacterial biofilm formation, we first examined biofilm biomass and matrix development. While a mature *P. aeruginosa* biofilm with bacterial cells clearly embedded within an EPS matrix developed on pNGPDA over 24 h, this was not the case on pEGdPEA where only a few cells and cell clusters lacking matrix were apparent. After 72 h the biofilm biomass on pNGPDA increased >2-fold while no biofilm formed on pEGdPEA. On a 1:1 p(NGPDA-co-EGdPEA) copolymer surface, biofilm formation was substantially less than on pNGPDA after 24 h and in contrast to pNGPDA, the biofilm was dispersed after 72 h incubation. This demonstrated that pEGdPEA component of the copolymer was capable of inhibiting the biofilm promoting properties of the pNGPDA component. These data suggested that early interactions with pEGdPEA were most likely key to biofilm inhibition.

*P. aeruginosa* explores surfaces via flagellar- and TP4-mediated motility (de Anda, Lee et al. 2017). Since near surface single cell tracking via DHM did not reveal any differences in swimming speed above pNGPDA compared with pEGdPEA, the behaviour of cells after surface contact was examined. Early-stage reversible attachment of *P. aeruginosa* cells on a surface can be processive or non-processive depending on whether cells commit to stay on the surface long enough to either divide or detach before dividing (Lee, Vachier et al. 2020). Monotrichous *P. aeruginosa* cells also undergo surface-induced asymmetric cell division whereby a flagellated parent cell gives rise to a piliated daughter that stays on the surface and engages in local T4P-mediated surface exploration while the parent detaches from the surface and swims away (Laventie, Sangermani et al. 2019). While no differences were observed during the first hour following inoculation onto the surface, over the next 2 h, greater numbers of bacteria were observed to accumulate and remain for longer periods on pNGPDA than on pEGdPEA. Residence times on pNGPDA were noticeably much more heterogeneous than on pEGdPEA. We also found that after cell division for one-legged branching, 44% left pEGdPEA compared with 17% for pNGPDA suggesting that surface chemistry impacts on asymmetric cell division and influences the decision of *P. aeruginosa* cells to stay or to leave. Of the cells which remained on the two surfaces, no major differences in the proportion of motile cells, their directionality or surface orientation were apparent. Given that Psl acts as both an adhesive (Ma, Jackson et al. 2006) and as a signal that stimulates c-di-GMP production (Irie, Borlee et al. 2012), these data are consistent with our observation that c-di-GMP levels increase on pNGPDA to a much greater extent than on pEGdPEA. During biofilm formation on borosilicate glass, the Wsp system generates heterogeneity in *P. aeruginosa* surface sensing, resulting in two physiologically distinct subpopulations of cells, one which is sessile with high levels of c-di-GMP and EPS and one which has low levels and is highly motile (Armbruster, Lee et al. 2019). We observed similar c-di-GMP heterogeneity on pNGPDA but not on pEGdPEA where c-di-GMP remained consistently low, in agreement with the lack of Psl trails. Our data showing reduced residency on pEGdPEA is also consistent with that obtained for a *P. aeruginosa pslD* mutant (which cannot produce Psl) on glass (Zhao, Tseng et al. 2013). Further confirmation that *P. aeruginosa* does not produce adhesive EPS on pEGdPEA was obtained from shear stress experiments which revealed that the force required to remove *P. aeruginosa* wild type cells was 27-fold lower than for pNGPDA.

The polar flagellum of *P. aeruginosa* has been proposed to serve as the primary surface sensor for both rapid activation of T4P assembly and stable surface attachment such that T4P subsequently act as independent surface sensors driving up production of c-di-GMP, EPS production and hence biofilm development (Laventie, Sangermani et al. 2019, Harrison, Almblad et al. 2020). To understand the relative contributions of flagellar and T4P to differential sensing of pNGPDA and pEGdPEA surfaces, we employed Δ*fliC* and Δ*pilA* mutants. By tracking the fate of single cells on the two polymer surfaces after cell division within 1-2 h after initial contact, we observed that for both mutants, 95% of the cells remained on the surface. While the Δ*pilA* mutant was unable to progress to biofilm formation on either polymer consistent with its weak adherence, the Δ*fliC* mutant adhered so strongly to both pNGPDA and pEGdPEA that the shear force required was greater than the maximum that could be applied. Mutations that disrupt flagellar assembly are known to stimulate adhesion and EPS production in bacteria including *P. aeruginosa* (Harrison, Almblad et al. 2020), *Vibrio cholerae* (Watnick, Lauriano et al. 2001) and *Caulobacter crescentus* (Hershey, Fiebig et al. 2020). However, despite the increased adhesion strength, the *P. aeruginosa* Δ*fliC* mutant formed less biofilm biomass on pNGPDA than the wild type but still behaved differentially on the two polymers as was unable to form a biofilm on pEGdPEA. These data indicate that strength of cell adhesion alone is likely insufficient to initiate biofilm formation on pEGdPEA and that the *P. aeruginosa* Δ*fliC* mutant, in common with the Δ*pilA* mutant wild type is unable to progress to the later stages of biofilm development on this polymer. Although a *P. aeruginosa* PAO1 Δ*pilA* Δ*fliC* double mutant did not show reduced Psl production after growth on VBMM agar (Harrison, Almblad et al. 2020), we found that a Δ*pilA* Δ*fliC* mutant was unable to form a biofilm on pNGPDA indicating that T4P are essential for biofilm formation under the conditions employed.

In contrast to the Δ*fliC* mutant, the *P. aeruginosa* Δ*motAB* Δ*motCD* double mutant which is non-motile but assembles a complete flagellum showed only weak adhesion to, and did not form a biofilm on either polymer, confirming the importance of a functional and powered flagellum for biofilm formation. However, the two stators are not functionally equivalent. MotCD appears to generate torque more efficiently than MotAB and is required for swimming under high-load conditions and for swarming (Baker and O’Toole 2017) whereas *motAB* is involved in T4P assembly (Toutain, Caizza et al. 2007, Baker and O’Toole 2017) (Laventie, Sangermani et al. 2019). In addition, Schneiderbrand et al (2019) observed that in tethered bacteria (Schniederberend, Williams et al. 2019), MotCD is required to maintain flagellum rotation while MotAB stops the flagellum rotating consistent with a role for MotAB in supporting surface residence via the deployment of T4P. On pEGdPEA and pNGPDA, neither MotAB nor MotCD facilitated differentiation between the two polymers when the early stage post division fate of the cells was considered during the first 1-2 h following surface engagement. However, a far greater number of *motAB* mutant cells departed from either surface compared with the *motCD* cells suggesting that MotCD is key to early-stage surface departure from both polymers. This is consistent with the ability of MotCD to maintain the flagellum rotation necessary for cells to detach from a surface (Schniederberend, Williams et al. 2019). In contrast, the greater number of *motCD* mutants remaining on the surface reflects the contribution of MotAB to irreversible attachment (Toutain, Caizza et al. 2007). This is also consistent with the greater adhesion strength and higher biomass for the *motCD* mutant compared with the *motAB* mutant although both were substantially lower than wild type on pNGPDA.

In *P. aeruginosa* strain PA14, a transposon mutant exhibiting a defect in biofilm formation and incapable of transitioning from reversible to irreversible attachment was mapped to a gene termed *sadB* (Caiazza and O’Toole 2004, Caiazza, Merritt et al. 2007, Toutain, Caizza et al. 2007)(Caiazza and O’Toole 2004, Caiazza, Merritt et al. 2007, Toutain, Caizza et al. 2007). Overexpression of *sadB* enhanced biofilm production in a *mot* stator dependent manner (Caiazza and O’Toole 2004, Caiazza, Merritt et al. 2007, Toutain, Caizza et al. 2007). Furthermore, Harrison *et al* (2020) reported that the elevated Psl production in a PAO1 *fliC* mutant grown on an agar surface could be suppressed by both *sadB* and *motAB* mutations (Harrison, Almblad et al. 2020). The data presented here are consistent with these findings given that neither *sadB* nor *mot* mutants were able to form a biofilm on pNGPDA. These data suggest that on pEGdPEA, *P. aeruginosa* is unable to progress from reversible to irreversible attachment as it was unable to induce *sadB* expression. In common with c-di-GMP production, we observed that the *sadB* promoter was expressed heterogeneously at a much lower level on pEGdPEA than on pNGPDA and that constitutive expression of *sadB* from a multicopy plasmid substantially increased the strength of adhesion. This in turn enabled the *P. aeruginosa* wild type to overcome the biofilm resistance of pEGdPEA in a Mot flagellar stator dependent manner consistent with the increase in cdiGMP production in the wild type but not *motABCD* mutant constitutively expressing s*adB*. These findings suggested that inhibition or lack of activation of SadB blocks biofilm formation on pEGdPEA by controlling the transition from reversible to irreversible attachment. Although the biochemical function of the *P. aeruginosa* SadB protein has yet to be elucidated, a *Pseudomonas fluorescens* SadB orthologue has been reported to bind c-di-GMP (Muriel, Blanco-Romero et al. 2019). In *P. aeruginosa* strain PA14, SadB functions downstream of the SadC/BifA/c-di-GMP regulatory pathway and is involved in regulating the Pel expolysaccharide (Caiazza and O’Toole 2004, Merritt, Brothers et al. 2007).

These findings demonstrate for the first time, a unique mechanism by which a simple polymer prevents bacterial biofilm formation by inhibiting the transition from reversible to irreversible attachment. Further work will be required to confirm the biochemical function of SadB *in P. aeruginosa* and the precise molecular mechanism that results in the inhibition of *sadB* expression on pEGdPEA and other biofilm resistant acrylate polymers. They also highlight the potential of such materials for preventing biofilm formation in diverse healthcare, industrial and natural environments.

## Methods

### Bacterial strains and culture conditions

**Table S3** lists the bacterial strains and plasmids used. *P. aeruginosa* strain PAO1-W (University of Washington subline) and *E. coli* strains were routinely cultured in lysogeny broth (LB) at 37°C with shaking (200 rpm). Where required the following antibiotics were added: ampicillin (Ap) 100 µg.ml^-1^ (*E. coli*); carbenicillin (300 µg.ml^-1^) (*P. aeruginosa*); tetracycline (Tc) 25 µg.ml^-1^ (*E. coli*) or 125 µg.ml (*P. aeruginosa*).

### *P. aeruginosa* deletion mutants, complementation plasmids and reporter gene fusions

*P. aeruginosa* Δ*pilA,* Δ*fliC,* Δ*pilA*Δ*fliC* and Δ*sadB* deletion mutants were constructed via allelic exchange. Two PCR products amplifying the upstream and the downstream regions of each gene were generated using the primer pairs 1FW/1RW, and 2FW/2RW respectively (**Table S4**). The resulting PCR products were cloned into the suicide plasmid pME3087 (Voisard, Bull et al. 1994) and introduced into *P. aeruginosa* via conjugation with *E. coli* S17-1 λpir. Recombinants were selected on tetracycline followed by enrichment on carbenicillin (Malone, Jaeger et al. 2012). Deletions were confirmed by DNA sequence analysis and their phenotypes confirmed (**Fig. S9**). For genetic complementation of the corresponding *P. aeruginosa* mutants, the Δ*pilA,* Δ*fliC* and Δ*sadB* genes were PCR-amplified using chromosomal DNA as a template and cloned into the shuttle vector pME6032 which contains a *lacIQ* mutation rendering the P*tac* promoter constitutive (Heeb, Blumer et al. 2002, Popat, Crusz et al. 2012). The *sadB*’::mNeonGreen transcriptional reporter fusion was constructed by amplifying the mNeonGreen gene by PCR from pNCS-mNeonGreen (Allele Biotechnology) using the primer pair mNeonGFW/mNeonGRV (**Table S4**) and ligated as a SphI-PstI fragment with pUCP22Not, resulting in pUCP22Not-*mNeonGreen*-T0-T1 (pMP5). A 400bp-PCR product amplifying the region upstream of the *sadB* gene using the primer pair PsadBFW/PsadBRV-RBSII was inserted into pMP5 at the XbaI/SphI sites, generating pUCP22Not-P*_sadB_*-RBSII-*mNeonGreen*-T0-T1 (pMP6).

### Polymer preparation and coating

All synthesis reagents were obtained from Sigma-Aldrich. Droplets of EGdPEA and NGPDA (5 µL) mixed with 1% (w/v) of photo-initiator (2,2-dimethoxy-2-phenylacetophenone) were polymerized on glass cover slips or borosilicate glass slides by exposure of the liquid to UV photoinitiation under an argon atmosphere with less than 2000 ppm of oxygen. To avoid delamination, methacrylate silaninsation of the glass after oxygen plasma cleaning was employed. The same approach was used to polymerize BnMA, iBMA and TMCHMA.

### Polymer surface physico-chemical characterization

Surface roughness was determined using an atomic force microscope (Bruker Dimension Icon®, BruckerNano, Santa Barbara, US) in PeakForce QNM® mode with Bruker MSNL-F tips. Root-mean-square (rms) values for roughness were calculated using an average of at least five measurements over areas of 0.5 µm x 0.5 µm. Water contact angle (WCA) was measured using the sessile drop method on an automated Krüss DSA 100 instrument. Zeta potentials for pNGPDA and pEGDPEA in water, phosphate buffered saline and RMPI were measured using a Zetasizer Nano ZS (Malvern, UK). Monodispersed melamine particles were used as tracer particles at concentrations of 10-4 % (w/v). ToF-SIMS spectra were obtained using a ToF-SIMS IV (IONTOF GmbH) instrument operated using a 25 kV Bi_3_^2+^ primary ion source exhibiting a pulsed target current of >0.3 pA. Samples were scanned at a pixel density of 100 pixels per mm, with 8 shots per pixel over a given area. A primary ion dose of 2.45 x 10^11^ ions per cm^2^ was used for each sample area ensuring static conditions were maintained throughout. Both positive and negative secondary ion spectra were collected (mass resolution of >7000 at m/z = 29), over an acquisition period of 15 scans. As the polymer samples were non-conductive, charge compensation was applied using a low energy (20 eV) electron floodgun. Secondary ion peaks were identified using a peak search tool (SurfaceLab 6).

### Biofilm formation and confocal microscopy

Biofilms were cultivated on borosilicate glass slides spotted in triplicate with the relevant polymer on coated glass or cover slips. Fluorescently tagged *P. aeruginosa* strains were inoculated into petri dishes containing the coated cover slips in RPMI-1640 and incubated at 37°C with shaking for 24 h. Slides were examined directly under a laser scanning confocal fluorescent Microscope (LSM2, Zeiss) using excitation wavelengths of 488 nm and 587nm for eGFP and mCherry respectively. Imaging was carried out using Zen 2011 imaging software (Zeiss). A total of 5 Z-stacked images were collected per polymer spot. Biomass was calculated using Image J (NIH, Bethsda, MD, USA) and Comstat 2.1. Software package (www.cosmtat.dk, Lyngby, Denmark). eDNA in biofilms was stained with propidium iodide (0.6 μM).

Biofilms were also analysed using environmental scanning electron microscopy (ESEM), enabling the semi-hydrated samples to be images without a conductive coating to gain insight into their native state. After incubation with *P. aeruginosa*, polymer coated coverslips were adhered to sample holders using die-cut carbon conductive adhesive discs (SPI Supplies/Structure Probe, Inc., West Chester, PA, USA) and imaged on a Quanta 200FEG SEM microscope (FEI Company, Eindhoven, the Netherlands). Chamber parameters were equilibrated to <-10 °C and 238 Pa at 5-15 kV to progressively sublimate sample water content. Images (at least 20 per sample) were captured at 10,000× magnification.

### Multimode microscopy and cell tracking

Time-lapse imaging of single bacterial cells were captured using the bespoke multimode microscope described by (Hook, Flewellen et al. 2019). Samples were analysed at 37°C using a Nikon Eclipse Ti inverted microscope using a 40× N.A. = 1.4, W.D. = 0.17 mm oil objective and fitted with an environmental chamber (Okolab). Images were acquired every 2 min with an Orca-Flash 4.0 digital CMOS camera (Hamamatsu). In-line digital holographic imaging (DHM) employed a 685-nm LX laser (Obis). Differential interference contrast (DIC) brightfield and widefield epifluorescence imaging was achieved using a single channel white MonoLED light source (Cairn Research). Experiments were conducted using a custom designed coverslip holder. Single bacterial cells were tracked using the bespoke MATLAB scripts described in (Hook, Flewellen et al. 2019). Instantaneous speed was calculated as a function of the displacement vector as shown in Equation 1 where Δ*t* = *t_i_*_+1_ – *t_i_* while average speed was calculated according to Equation 2 where n is number of points in the track.

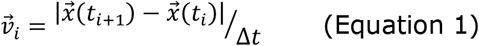

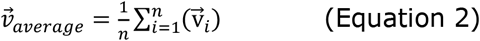

The radius of curvature *R* from centre of mass was calculated as described in Equation 3, where *R_i_* is the position vector of the i^th^ point on the trajectory and *R_cm_* is the centre-of-mass of all points as used before (Utada, Bennett et al. 2014).

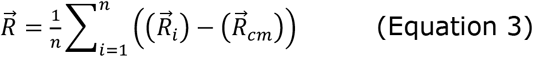

Bacterial cell residence times were measured as the time between the start and the end of tracks between t = 1 h to t = 2 h from inoculation. Cell division events were captured by searching for mother cells in frame *i* which divided into two daughter cells in frame *i^+1^*. These events (n=100 for each surface and bacterium over 3 independent replicates N=3) were manually assessed to avoid possible misreading of the automatic algorithm. Each daughter cell was assigned a new identification. The states of post-division cells were defined as follows with three types of division branching. “Two-legged” (d=2) branching occurs when both daughter cells detach, “one-legged” (d=1) when one daughter cell detaches, and “zero-legged” (d=0) when both daughter cells remain surface-attached.

To evaluate the distribution of T4P-mediated ‘crawling’ and ‘walking’ cells, the method of Rodesney *et al*. (Rodesney, Roman et al. 2017) was adapted. As a proxy readout for tilting, the projected aspect ratio in each frame was considered. A newly divided cell lying flat on the surface has an aspect ratio of ∼2. Therefore, a projected aspect ratio <2 indicates unambiguously that a cell is tilting. For each frame, cells were computed with either 1 (tilting up) or 0 (lying flat). For *P. aeruginosa* cells on pNGPDA and pEGdPEA, the average tilting fraction was considered as the average of the tilting fractions of cells over the duration of the experiment.

EPS trails on the polymer surfaces were visualised using Concanavalin A, Alexa Fluor™ 633 Conjugate (50 µg ml^-1^; ThermoFisher Scientific). *P. aeruginosa* cells were pipetted onto each surface and imaged in DIC mode for 2.5 h prior to further analysis by confocal microscopy. EPS trails were quantified as fluorescence intensity *F* using Zen 2011 imaging software (Zeiss) following Equation 4 where *Fb* and *Ft* are the intensities measured over an area of 100 µm^2^ on polymers at t = 0 h (background) and t = 2.5 h respectively.

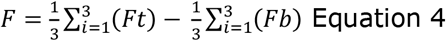

The transcriptional activities of *cdrA* and *sadB* promoters in *P. aeruginosa* on pEGdPEA and pNGPDA were quantified by measuring the fluorescence output from cells expressing *cdrA’::gfp*(ASV)c (**Table S3**) or *sadB’*::mNeonGreen (**Table S3**) over the first 3h after bacterial attachment. Tracks were obtained from images acquired using total internal reflectance fluorescence (TIRF) imaging every 30 s for 3h.

### Bacterial cell adhesion assay

A microfluidics apparatus based upon that described by Arpa-Sancet *et al*. (Arpa-Sancet, Christophis et al. 2012) was used to determine the adhesion strength of *P. aeruginosa* cells for each polymer. A flow chamber was constructed using xurography to create microchannels (14 mm×1.6 mm× 50 µm) by embedding double-sided silicone adhesive tape (50 µm; 3M™, USA) between a PDMS membrane and a polymer surface (Bartholomeusz, Boutte et al. 2005, Khashayar, Amoabediny et al. 2017). A computer-controlled low pressure precision pump (neMESYS Cetoni GmbH, Germany) was used to aspire medium and generate flow inside the channel. The system was mounted on an inverted microscope. Bacterial cells in RPMI at 37 °C were inoculated into the flow chamber for 10 min prior to applying an initial flow rate of 0.028 ml min^-1^ and a shear stress of ≈ 0.1 Pa. This was increased incrementally every 5 s until a maximum shear stress of ≈ 600 Pa was reached. Images were captured every 5 s and the shear stress (*τ*_50_) necessary to detach 50% of the surface-associated cells calculated. The shear stress (*τ*) is described as the force exerted onto channel walls which can be calculated using 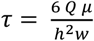 where Q is the flow rate, h and w are the height and the width of the channel respectively and µ is the liquid viscosity.

### Statistical analysis

Differences between data sets were assessed using unpaired *t* tests or one-way analysis of variance (ANOVA) as appropriate to determine the *P* value. The value *n* indicates the number of biological replicates. Error bars indicate ±1 standard deviation unit or standard error of mean as appropriate.

## Supporting information

SuppInformation

## Acknowledgments

This work was supported by Wellcome Trust joint senior investigator awards (Grant Nos. 103882 and 103884 to MRA and PW). AH was also supported by a University of Nottingham Research Fellowship. We thank N. Weston and for help with sample preparation and SEM (School of Life Sciences Imaging Facility), Alvin K. Teo for ToF-SIMS analysis and Francisco Pappalardo for AFM imaging.

## Author contributions

AC, JFD and ALH, MRA, PW conceived and designed the study. AC, PW, MRA, JFD, and ALH contributed to interpretation of data. JFD and MP constructed the mutants, reporters and conducted biofilm assays. OS and NF performed surface Z-potential measurements and DHM experiments respectively. AC and PW drafted the manuscript and all authors commented on the draft manuscript and approved submission.

