## Supplementary material for "Polymer-directed inhibition of reversible to irreversible attachment prevents *Pseudomonas aeruginosa* biofilm formation": SuppInformation

**Table S1.** Names, abbreviations and chemical structures of monomers used in this work.

| Monomer | Abbreviation | Chemical structure |
| --- | --- | --- |
| Ethylene glycol dicyclopentenyl ether acrylate | EGdPEA | 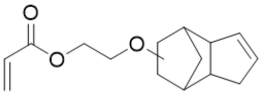 |
| Neopentyl glycol propoxylate diacrylate | NGPDA | 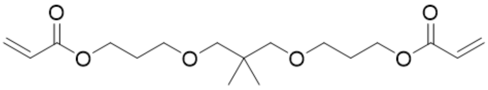 |
| Isobornyl methacrylate | iBMA | 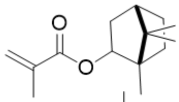 |
| Trimethylcyclohexyl methacrylate | TMCHMA | 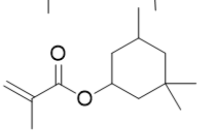 |
| Benzyl methacrylate | BnMA | 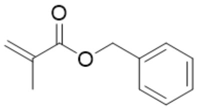 |

**Table S2.** Proportion of the population for each bacterial strain in different branching divisions considering pNGPDA (blue) and pEGdPEA (pink) between t = 1 h and t = 2 h.

| | d=0 | | d=1 | | d=2 | |
|---|---|---|---|---|---|---|
| | pNGPDA | pEGdPEA | pNGPDA | pEGdPEA | pNGPDA | pEGdPEA |
| PA wt | 83 ± 1 | 53 ± 10 | 17 ± 1 | 44 ± 7 | 0 ± 0 | 3 ± 3 |
| Δ*pilA* | 95 ± 1 | 95 ± 1 | 5 ± 1 | 5 ± 1 | 0 ± 0 | 0 ± 0 |
| Δ*fliC* | 95 ± 1 | 95 ± 1 | 5 ± 1 | 5 ± 1 | 0 ± 0 | 0 ± 0 |
| Δ*motAB* | 19 ± 4 | 21 ± 3 | 44 ± 7 | 54 ± 9 | 36 ± 5 | 24 ± 11 |
| Δ*motCD* | 53 ± 13 | 47 ± 4 | 54 ± 5 | 42 ± 5 | 12 ± 4 | 11 ± 1 |

**Table S3:** Bacterial strains and plasmids used in this study.

| Strain, plasmid or primers | Genotype and/or Relevant characteristic | Source or reference |
|---|---|---|
| **Strain** | | |
| ***P. aeruginosa*** | | |
| **PAO1-W** | Washington collection wild-type *P. aeruginosa* strain | University of Washington collection |
| **PAJD431** | In frame deletion of *pilA* in PAO1-W | (Carabelli, Isgró et al. 2020) |
| **PAJD477** | In frame deletion of *fliC* in PAO1-W | This study |
| **PAJD457** | In frame deletion of *fliC* in PAJD431 resulting in a Δ*pilA* Δ*fliC* in PAO1-W | This study |
| **PAJD360** | In frame deletion of *motAB* in PAO1-W | (Hook, Flewellen et al. 2019) |
| **PAJD552** | In frame deletion of *motCD* in PAO1-W | (Hook, Flewellen et al. 2019) |
| **PAJD549** | In frame deletion of *motCD* in PAJD360 resulting in a quadruple mutant Δ*motAB*Δ*motCD* in PAO1-W | (Hook, Flewellen et al. 2019) |
| **PAJD544** | In frame deletion of Δ*sadB* in PAO1-W | This study |
| ***E. coli*** | | |
| **DH5α** | *recA1 endA1 hsdR17 supE44 thi-1 gyrA96 relA1 Δ(lacZYA-argF)U169[ϕ80 dlacZΔM15],* Nal$^R$ | (Hanahan 1983) |
| **S17.1λpir** | *thi pro hsdR hsdM$^+$ recA RP4-2-Tc::Mu-Km::Tn7 λpir*, Gm$^R$ | (Simon, Priefer et al. 1983) |
| **Plasmids** | | |
| **pME3087** | Suicide vector for homologous recombination, ColE1 replicon, Mob; Tc$^R$ | (Voisard, Bull et al. 1994) |
| **pMRE147** | pPROBE-based vector harbouring mClover3; Cm$^R$, Gent$^R$ | (Schlechter, Jun et al. 2018) |
| **pNCS-mNeonGreen** | pNCS cloning vector carrying mNeonGreen; Amp$^R$. | Allele Biotechnology |
| **pME6032** | pVS1-p15A shuttle expression vector; IPTG inducible; Tc$^R$ | (Heeb, Blumer et al. 2002) |
| **pME6032Δ*lacIQ*** | pVS1-p15A shuttle expression vector containing a *lacIQ* mutation; Gent$^R$ | (Popat, Crusz et al. 2012) |
| **pMMR** | pME6032Δ*lacIQ* derivative containing constitutively expressed mCherry *orf*; Gent$^R$. | (Popat, Crusz et al. 2012) |
| **pCdrA::*gfp*(ASV)C** | pUCP22Not-$P_{cdrA}$-RBSII-*gfp*(ASV)-T0-T1; Amp$^R$, Gent$^R$ | (Rybtke, Borlee et al. 2012) |
| **pMP5** | pUCP22Not-*mNeonGreen*-T0-T1; Amp$^R$, Gent$^R$. | This study |
| **pMP6** | pUCP22Not-$P_{sadB}$-RBSII-*mNeonGreen*-T0-T1; Amp$^R$, Gent$^R$. | This study |
| **pJD112** | pME3087 derivative for the generation of *pilA* in frame deletion mutant; Tc$^R$. | (Becher and Schweizer 2000) |

| | | |
|---|---|---|
| **pJD113** | pME3087 derivative for the generation of *fliC* in frame deletion mutant; $Tc^R$. | This study |
| **pPilA** | pME6032 Δ*lacIQ* derivative containing a PCR fragment of 0,450 kbp with *pilA* gene; used for complementation; $Tc^R$. | (Heeb, Blumer et al. 2002) |
| **pFliC** | pME6032 Δ*lacIQ* derivative containing a PCR fragment of 1,467 kbp with *fliC* gene; used for complementation; $Tc^R$. | This study |
| **pSadB** | pME6032 Δ*lacIQ* derivative containing a PCR fragment of 1,410 kbp with *sadB* gene; used for complementation; $Tc^R$. | This study |

**Table S4** Oligonucleotide Primers used in this work.

| Name | Sequence 5′ to 3′ | Modification 5′-3′ |
|---|---|---|
| PilAΔFW1 | ATATCTAGAATGCCGAACTGCTCG | XbaI |
| PilAΔRV1 | TTAGTTATCACAACCTTGAGCTTTCATGAATCTCTC | N/A |
| PilAΔFW2 | TTCATGAAAGCTCAAGGTTGTGATAACTAAGGTGAT | N/A |
| PilAΔRV2 | TATCTGCAGAAGTGGAAGTGGAGA | PstI |
| FliCΔFW1 | ATATCTAGAATGCTCGAAGGCGCGCATCT | XbaI |
| FliCΔRV1 | TTAGCGCAGCAGGCTTGTAAGGGCCATGGTGATTTC | N/A |
| FliCΔFW2 | ACCATGGCCCTTACAAGCCTGCTGCGCTAAGCCCGG | N/A |
| FliCΔRV2 | TATAAGCTTAAGTCGTTCAACCCGCGCGT | HindIII |
| SadBΔFW1 | ATATCTAGAAGCGCCGCCAGCCACTGCGT | XbaI |
| SadBΔRV1 | TCACCCCGGCAAGCGGGCTTCTGTCATGACGAGACC | N/A |
| SadBΔFW2 | GTCATGACAGAAGCCCGCTTGCCGGGGTGACCGGGT | N/A |
| SadBΔRV2 | TATAAGCTTACCTGGTGTTCCAGTTGCAT | HindIII |
| PilAFW | TATGAATTCATGAAAGCTCAAAAAGGC | EcoRI |
| PilARV | TATCTCGAGTTAGTTATCACAACCTTT | XhoI |
| FliCFW | TATGAATTCATGGCCCTTACAGTCAAC | EcoRI |
| FliCRV | TATCTCGAGTTAGCGCAGCAGGCTCAG | XhoI |
| SadBFW | TATGAATTCATGACAGAAGCCGCCCTG | EcoRI |
| SadBRV | TATCTCGAGTCACCCCGGCAAGCGATT | XhoI |
| mNeonGFW | AAAGCATGATGAATTCGAAAGGCGAAGA | SphI |
| mNeonGRV | AAACTGCAGCTTATTTATATAATTCATCCATTCCCATAAC | PstI |
| PsadBFW | AAATCTAGACCACTGCGTGCTGCTCTAC | XbaI |
| PsadBRV-RBSII | TTTGCATGCTCATAGTTAATTTCTCCTCTTTGACGAGACC ATGCGAGCTGG | SphI |

*nucleotide sequences of restriction sites are underlined.

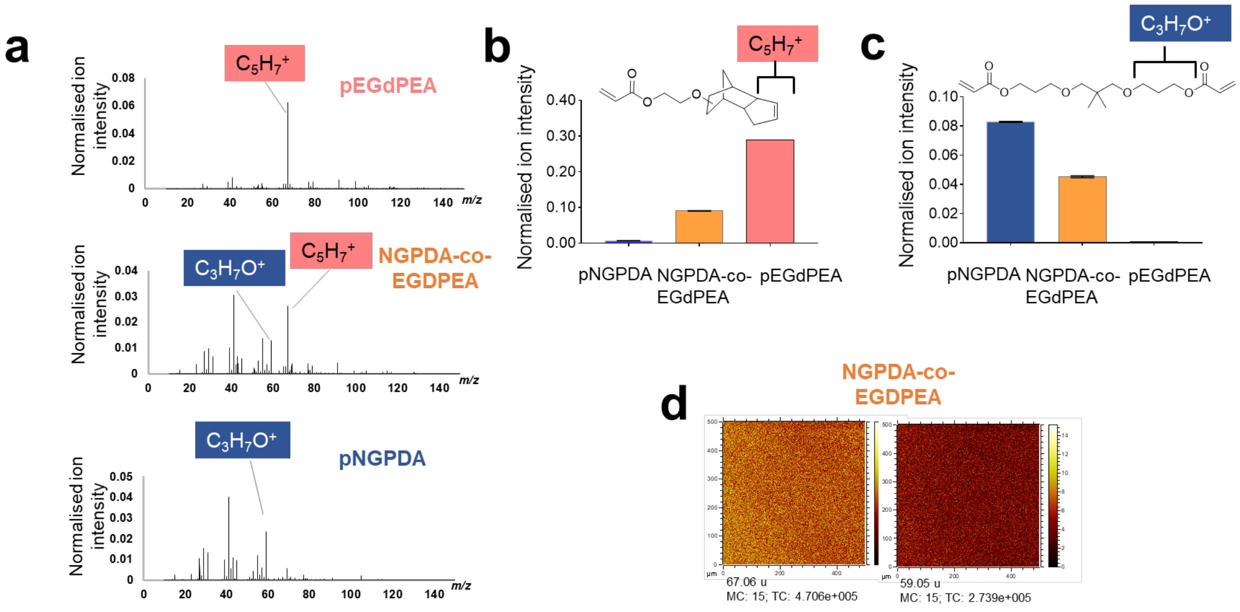


**Figure S1.** Polymer Surface characterisation. (**a**) ToF-SIMS characterization of polymer-coated coverslips. Positive ion ToF-SIMS spectra of pEGdPEA (pink), NGPDA-co-EGdPEA (1:1) (orange), pNGPDA (blue). $C_5H_7^+$ (*m/z* 67.06) and $C_3H_7O^+$ (*m/z* 59.05) are characteristic ions for pEGdPEA and pNGPDA respectively; comparison of normalised ion intensities for $C_5H_7^+$ (**b**) and $C_3H_7O^+$ (**c**) for the 3 polymers; (**d**) An example corresponding to a secondary ion image (500 x 500 µm) for both $C_5H_7^+$ (m/z 67.06) and $C_3H_7O^+$ (m/z 59.05) ions using pNGPDA-co-EGdPEA. The two monomers polymerised homogeneously.

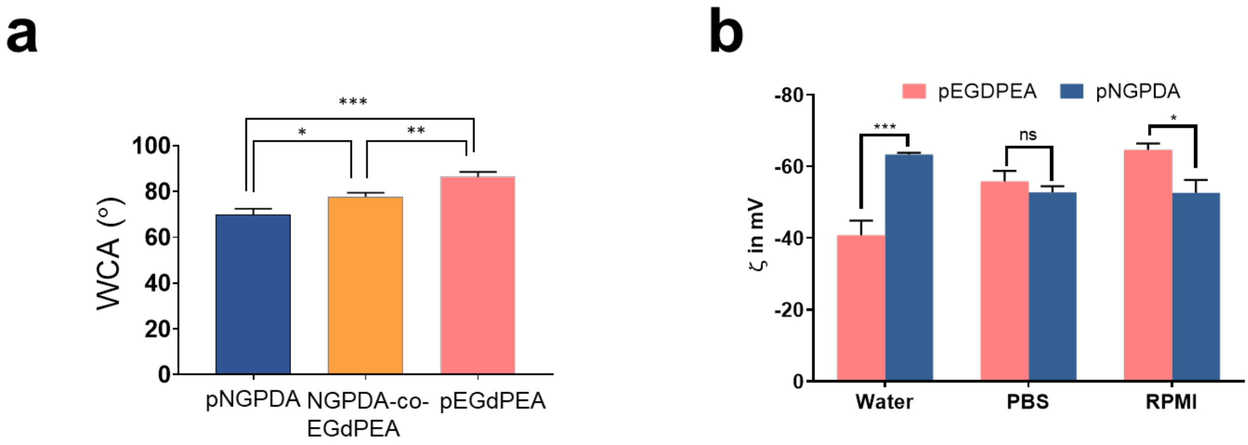


**Figure S2.** (**a**) Water contact angle (WCA) measurements on pNGPDA (blue), pEGdPEA (pink) and pNGPDA-co-EGdPEA (orange). Error bars show ± 1 standard deviation (N=9). ***$p$<0.001, **$p$< 0.01 and *$p$<0.05. Statistics assessed using One-way ANOVA with Tukey's multiple correction. (**b**) Surface zeta potential measured on films of pNGPDA and pEGDPEA in water, phosphate buffered saline (PBS) and RPMI. $p$ values < 0.05. Error bars show mean ± 1 s.d (n=2). Significance was determined by paired t-test comparisons. ***$p$<0.001 and *$p$<0.05.

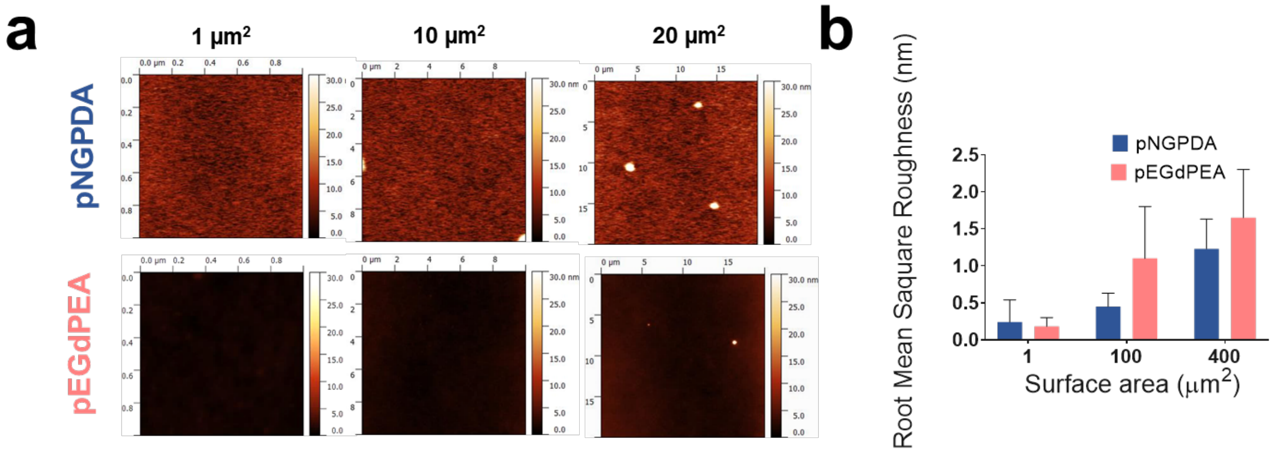


**Figure S3.** (**a**) Atomic force microscopy height images of pNGPDA (blue) and pEGdPEA (pink) for areas of 1×1, 10×10 and 20×20 µm$^2$. Images were previously line flattened to remove surface curvature and slope; (**b**) Determination of the root mean square roughness (nm) of the polymers at 1 µm$^2$, 100 µm$^2$ and 400 µm$^2$ from the surfaces in dry conditions. Values are the mean of four images taken over 4 different samples. The error bars equal ± 1 standard deviation (N=4). Significance was determined by analysis of variance 2-way ANOVA and Sidak's post-test comparison for differences between the samples.

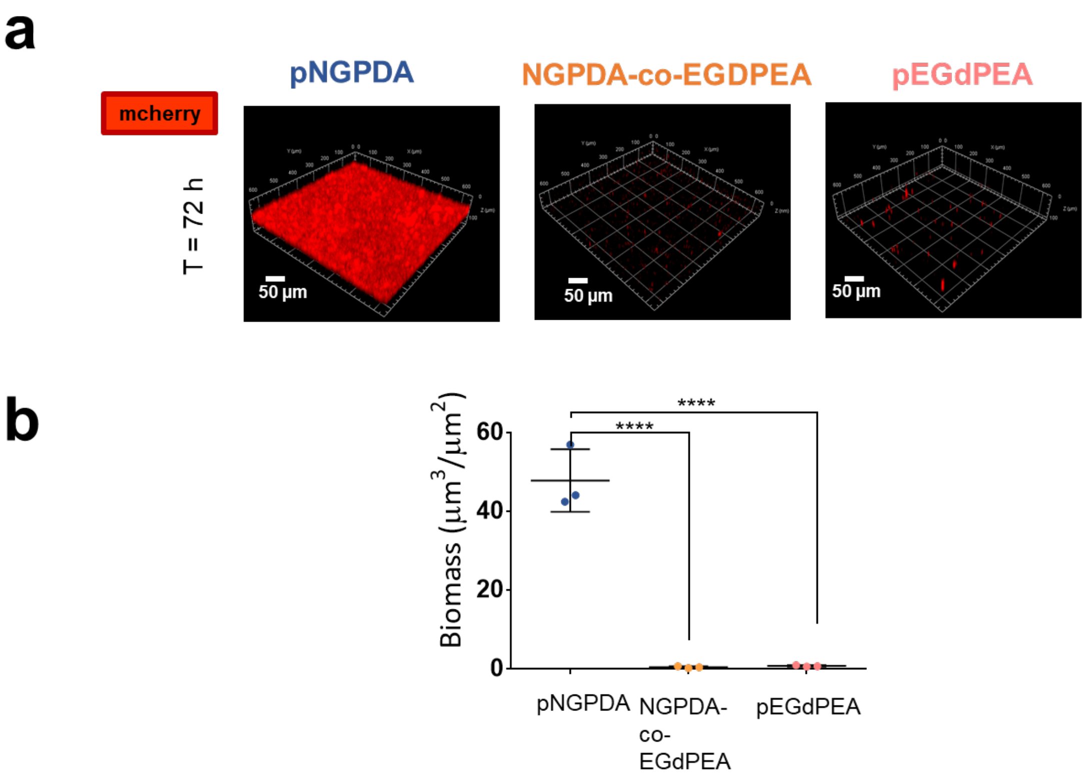


**Figure S4.** *P. aeruginosa* PAO1-W biofilm formation after 72 h on pNGPDA, pNGPDA-co-EGDPEA and pEGdPEA (**a**) Confocal images (10×, 0.3) of *P. aeruginosa m-cherry* tagged after 24 h (top) and 72 h (bottom) on pNGPDA (left), pNGPDA-co-EGDPEA (middle), and pEGdPEA (right). Samples were washed 2× in PBS and 1× in H2O. Scale bar, 50 µm; (**b**) Quantification of bacterial biomass on the 3 polymer surfaces. Error bars show ± 1 standard deviation (N=3). Significance was determined by analysis of variance One-way ANOVA and Tukey's post-test comparison for differences between the indicated samples. **** $p$<0.001, *** $p$<0.001.

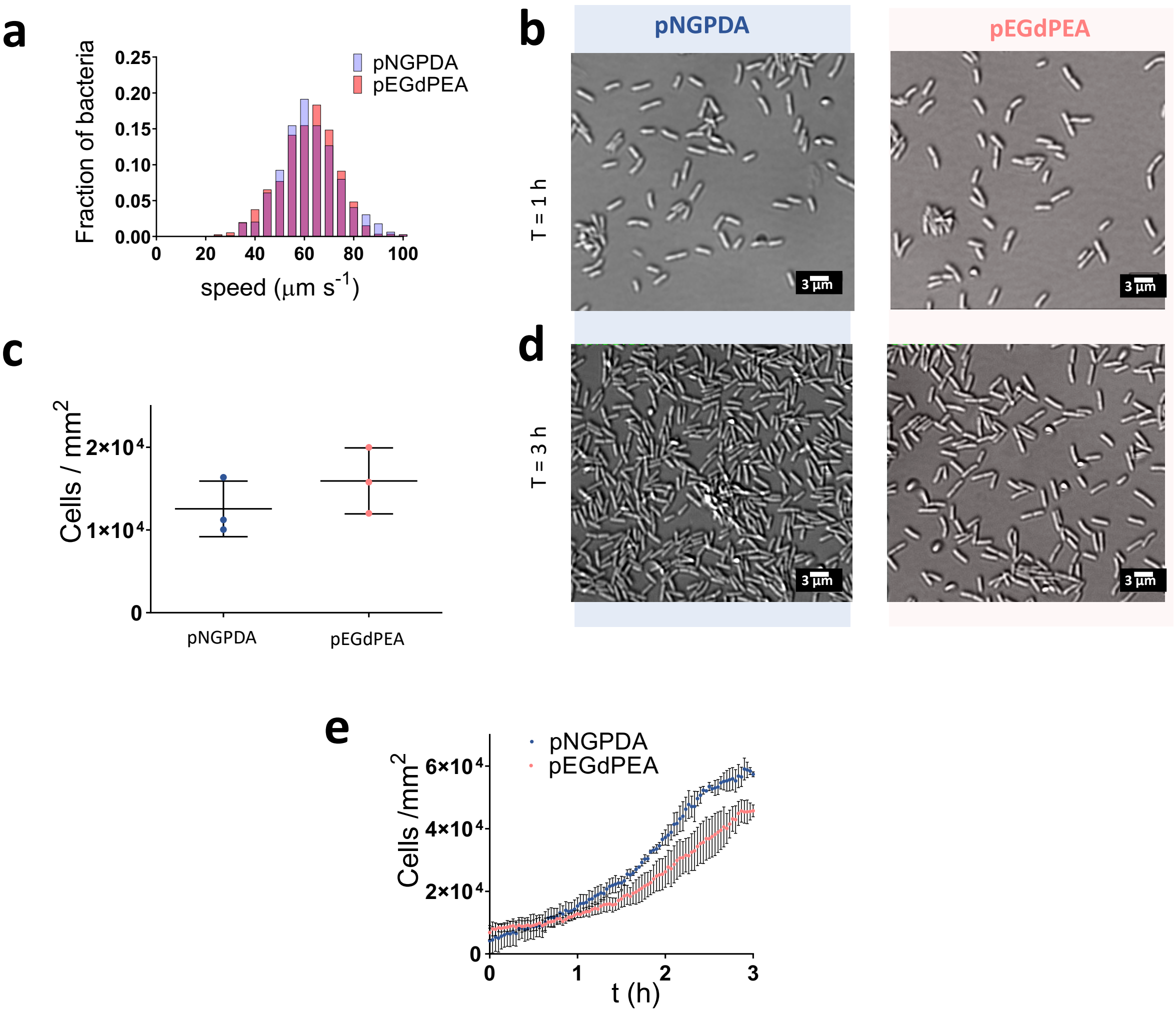


**Figure S5. Swimming motility above and accumulation on pNGPDA and pEGPDA.** Digital holographic microscopy (DHM) analysis up to 50 µm above each surface showing (**a**) average swimming speed of tracks above pNGPDA (blue) and pEGdPEA (pink) (N=3) considering n>600 tracks for each sample of log phase bacteria over a 10 min period immediately after inoculation into the chamber. Bin is 5 and 0.5. Differential interference (DIC) microscopy tracking (**b** and **d**) of cells at the surface showing images of PAO1 accumulating on pNGPDA and pEGdPEA after 1 h (**b**) and 3h (**d**) after inoculation; magnification, 40×; scale bar, 3 µm. (**c**) Categorical scatter plot of bacterial surface coverage (T = 1h). Error bars show ± 1 standard deviation (N=3). Scale bar 3 µm; (**e**) Number of bacteria per frame *vs* time increases faster for pNGPDA (blue) than pEGdPEA (pink). Dots represent mean ± 1 standard deviation (N=3).

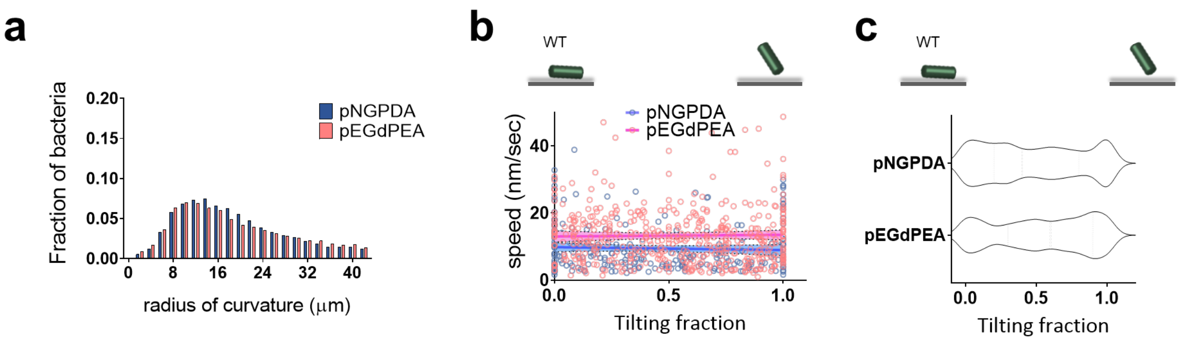


**Figure S6.** Directional motility and tilting of single *P. aeruginosa* cells (**a**) Histogram of radius of curvature for all tracks on different polymers. n>12,000 (N=3). A higher radius of curvature corresponds to more directional motility. Bin is 2. (**b**) Scatter plot showing the mean speed of cells during an entire trajectory as a function of the fraction of the cell's tracked lifetime that was spent tilting up from the *crawling* orientation into the vertical position (*walking*). Linear fits (y = m*x + b) to each dataset yield: for pEGdPEA, m = 0.5 ± 1.3, b = 13.0 ± 0.8; for pNGPDA, m = -0.9 ± 3.0, b = 9.8 ± 1.2 (c) Violin plot showing the proportion of cells as a function of tilting fraction.

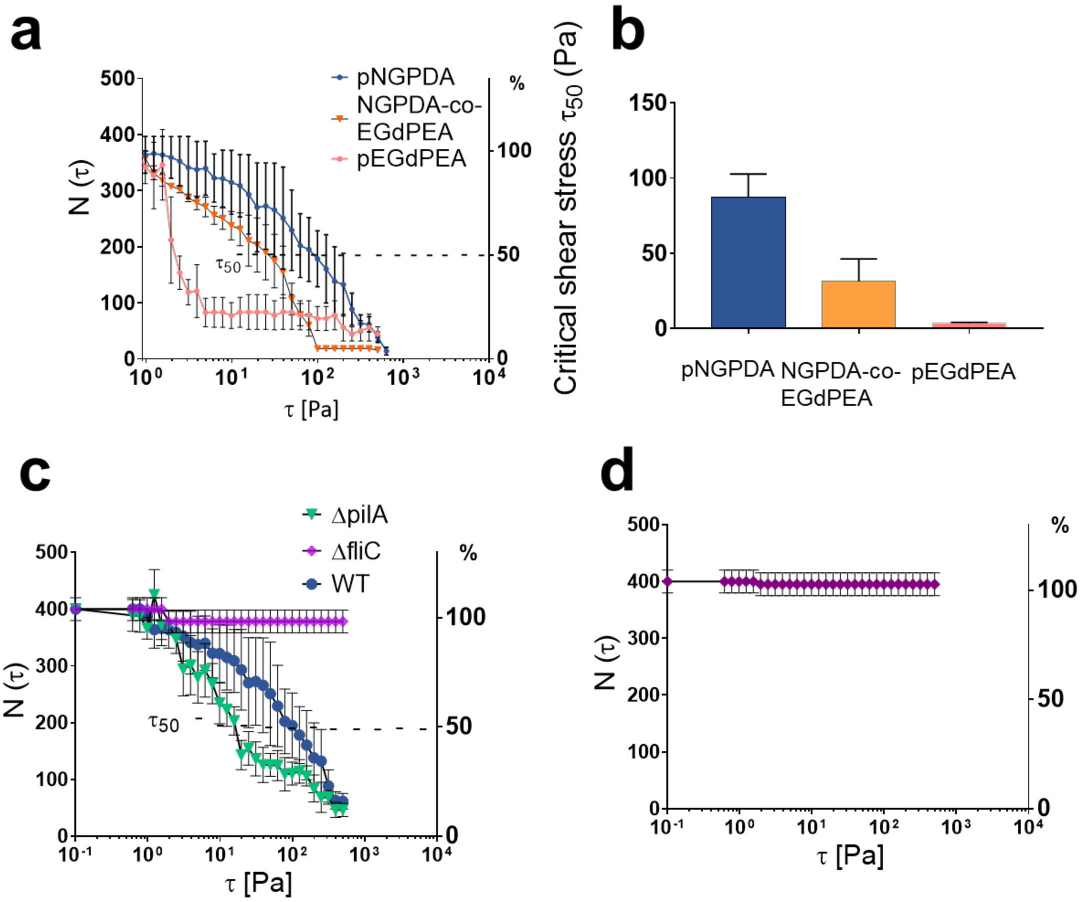


**Figure S7.** Measurement of *P. aeruginosa* PAO1-W wild type, flagella *(fliC)* and T4P *(pilA)* mutant adhesion strength on pNGPDA, pEGdPEA and pNGPDA-co-EGDPEA surfaces. (**a**) number of PAO1-W surface associated cells on pNGPDA (blue-circles), pEGdPEA (pink-circles) and pNGPDA-co-EGdPEA (orange-triangles) surfaces with increasing shear stress $\tau$ (Pa). Cells were incubated at the presence of the surface for 10 min.(**b**) Critical shear stress ($\tau_{50}$) necessary to detach 50% of PAO1-W cells and (**c**) number of *P. aeruginosa* Δ*pilA* cells (green), Δ*fliC* (purple) and PAO1-W (blue) on pNGPDA with increasing shear stress $\tau_{50}$ (Pa) and (**d**) Δ*fliC* on pEGdPEA. Error bars represent ± 1 SEM (N=3).

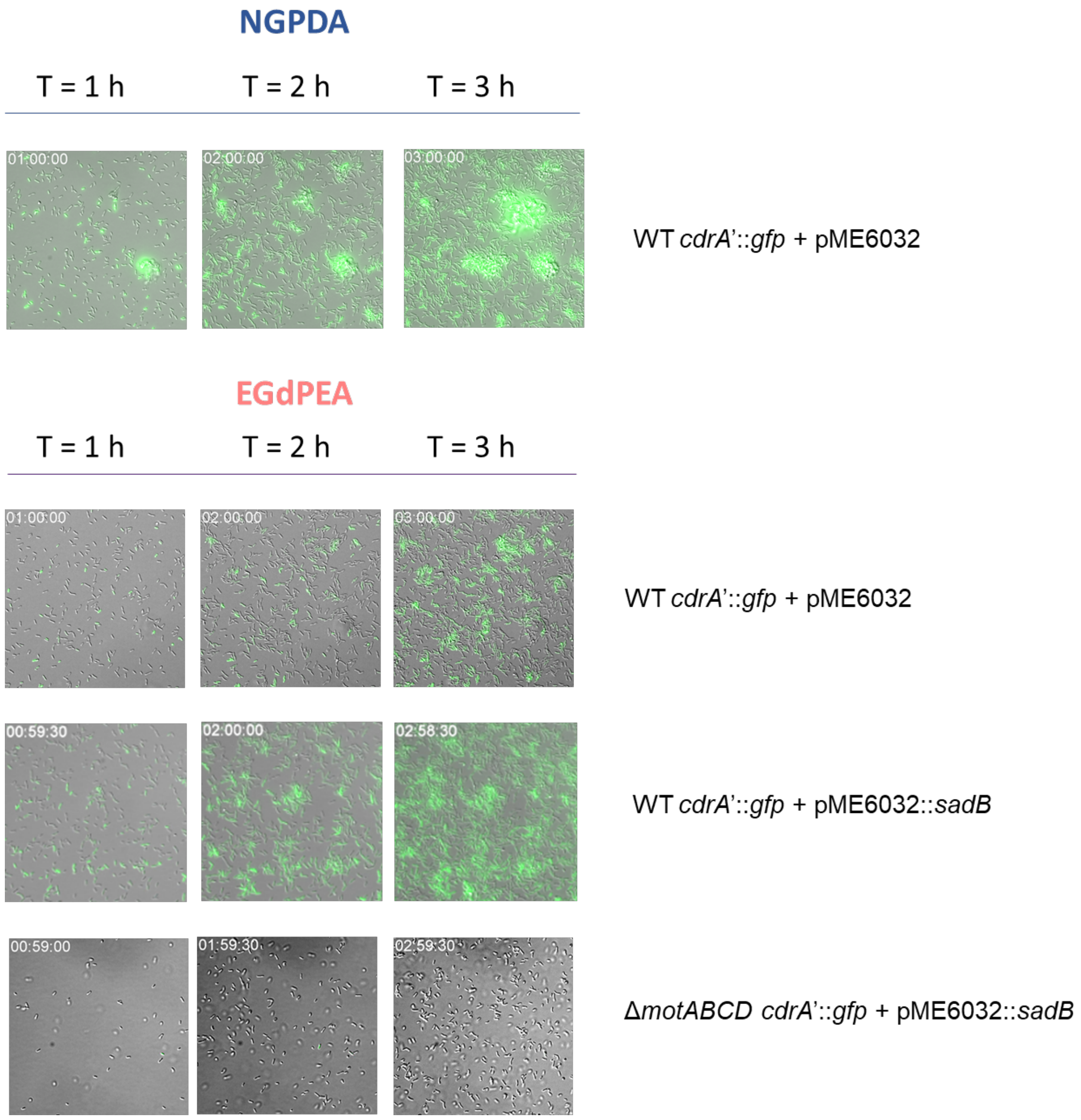


**Figure. S8** The impact of constitutive *sadB* on *cdrA'::gfp* expression in the *P. aeruginosa* wild type (WT) and Δ*motABCD* mutant on pEGdPEA. Representative epifluorescence microscopy images taken at t=1, 2 or 3 h after inoculation onto pNGPDA (wild type control) or pEGdPEA (WT transformed with backbone vector pME6032), WT transformed with both pSadB and pCdrA'::gfp(ASV) and ΔmotABCD mutant transformed with both pSadB and pCdrA'::gfp(ASV). Image size is 150 x 150 µm.

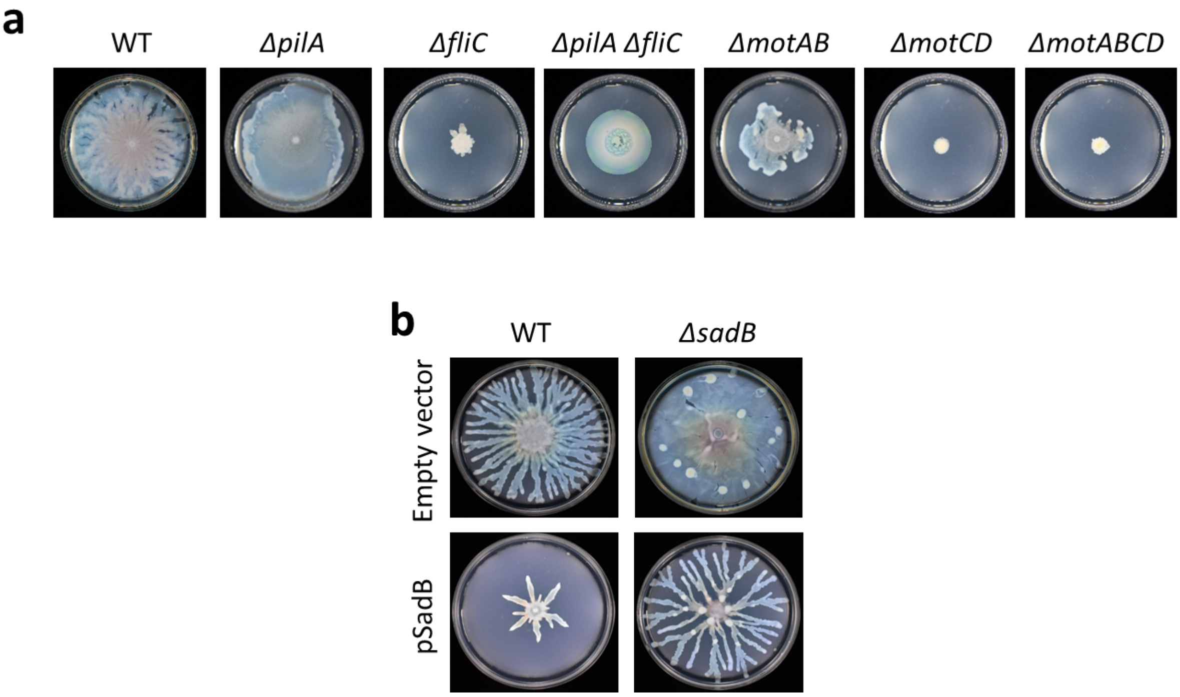


**Figure S9.** The swarming/sliding phenotypes of *P. aeruginosa* PAO1 wild type and the corresponding isogenic flagellar, flagellar stator, type IV pili, *sadB*, genetically complemented *sadB* mutant and the wild type strain constitutively expressing plasmid-borne *sadB* are shown. Each strain was plated on 0.5% swarming agar plates incubated overnight at 37°C for 16 h. (**a**) Representative swarming plates showing PAO1-W and Δ*pilA*, Δ*fliC*, Δ*pilA*Δ*fliC*, Δ*motAB*, Δ*motCD* and Δ*motABCD* mutant strains. (**b**) Representative swarming plates showing the PAO1-W and Δ*sadB* mutant transformed with the empty vector or a plasmid expressing *sadB*. The swarming/sliding phenotypes shown are consistent with those reported in the following references:
